# Impact of BA.1, BA.2, and BA.4/BA.5 Omicron Mutations on Therapeutic Monoclonal Antibodies

**DOI:** 10.1101/2022.12.25.521903

**Authors:** Bahaa Jawad, Puja Adhikari, Rudolf Podgornik, Wai-Yim Ching

## Abstract

The emergence of Omicron SARS-CoV-2 subvariants (BA.1, BA.2, BA.4, and BA.5) with an unprecedented number of mutations in their receptor-binding domain (RBD) of the spike-protein has fueled a new surge of COVID-19 infections, posing a major challenge to the efficacy of existing vaccines and monoclonal antibody (mAb) therapeutics. Here, a thorough and systematic molecular dynamics (MD) simulation study is conducted to investigate how the RBD mutations on these subvariants affect the interactions with broad mAbs including AstraZeneca (COV2-2196 and COV2-2130), Brii Biosciences (BRII-196), Celltrion (CT-P59), Eli Lilly (LY-CoV555 and LY-CoV016), Regeneron (REGN10933 and REGN10987), Vir Biotechnology (S309), and S2X259. Our results show a complete loss of binding for COV2-2196, BRII-196, CT-P59, and LY-CoV555 with all Omicron RBDs. REGN10987 also loses its binding against BA.1 but partially retains against BA.2 and BA.4/5. The reduction in binding is either significant for LY-CoV016 and REGN10933 or moderate for COV2-2130. S309 and S2X259 retain their binding strength against BA.1 but decrease against others. We introduce a mutational escape map for each mAb to identify the key RBD sites and critical mutation. Overall, our findings suggest that majority of therapeutic mAbs have diminished or lost their activity against Omicron subvariants, indicating the urgent need for a new therapeutic mAb, modifying current ones with a better mAb design, or seeking an alternative approach.

## 1. Introduction

Despite significant progress in understanding the biological implications of SARS-CoV-2 mutations, the emergence of several variants of concern (VOCs) poses the biggest new COVID-19 threat. These VOCs have different phenotypes from the original Wild-type (WT) strain in terms of infectivity, antigenicity, and transmissibility, resulting in increased infection and mortality rates globally.^1^ So far, the World Health Organization (WHO) has designated five VOCs: Alpha (B.1.1.7), Beta (B.1.351), Gamma (P.1), Delta (B.1.617.2), and Omicron (B.1.1.529 or BA.1). The latter one is the most mutated VOC with 37 mutations in the spike (S) protein, 15 of which are in the receptor-binding domain (RBD).^2^ This high number of RBD Omicron mutations represents a serious concern to the effectiveness of therapeutic monoclonal antibodies (mAbs), vaccines, and drugs.^3,4^ Another curious feature of Omicron is that it includes different subvariants (i.e., BA.1, BA.2, BA.3, BA.4, BA.5, etc.), which raises more doubts about the efficacy of these medical options. Indeed, many studies have demonstrated that Omicron subvariants can rise infectivity, reduce antibodies and vaccines effectivity, and result in more vaccine-breakthrough infections.^5–9^ Therefore, the study of the mutation effects of different Omicron sublineages is of paramount importance to predict which of them can help SARS-CoV-2 elude mAb therapies and vaccines. It may also enable us to prepare for future new variants.

RBD mutations in Omicron subvariants are either common or unique (**Figure 1**). Specifically, BA.2 has 16 RBD mutations, 12 of which are identical to those detected in BA.1 (G339D, S373P, S375F, K417N, N440K, S477N, T478K, E484A, Q493R, Q498R, N501Y, and Y505H), three of which are unique (T376A, D405N, and R408S), and the final one is S371F vs S371L of BA.1. But BA.2 does not have the G446S and G496S present in BA.1. On the other hand, both BA.4 and BA.5 (from here on referred to as BA.4/5) have accommodated the same 17 RBD mutations, 15 of which are similar to BA.2, with two additional mutations being L452R and F486V, but they lack Q493R harbored by BA.1 and BA.2. Strikingly, some of these RBD Omicron mutations have previously been identified in other VOCs, such as K417N, L452R, T478K, mutation at site 484, and N501Y, which lead to antibody escape and/or high infectiousness.^10^ However, many of Omicron mutations are extremely rare and their biological function remains to be determined. Consequently, there is an urgent need to identify their roles, particularly how they affect the immune recognition process of a wide range of clinical mAbs. This knowledge could provide critical information on the antigenic drift of RBD Omicron mutations, guiding the development of effective mAb therapeutics against SARS-CoV-2 variants.

**Figure 1.**
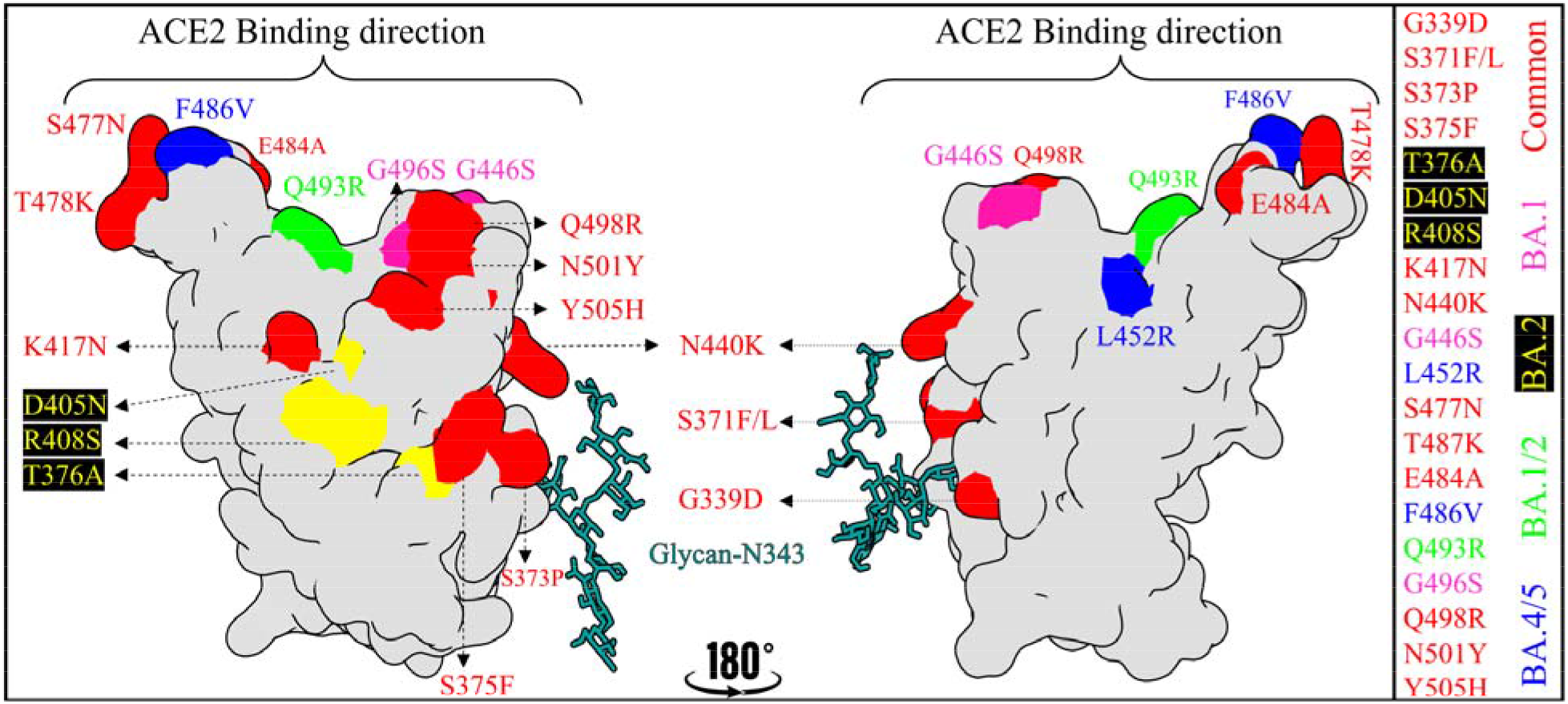
The RBD Omicron subvariant mutations are either common or unique. RBD shows in surface representation. The common RBD Omicron mutations across BA.1, BA.2 and BA.4/5 are marked and shown in red while the unique BA.1, BA.2 and BA.4/5 RBD mutations are represented by pink, yellow and blue respectively. Q493R mutation which only harbored by BA.1 and BA.2 is shown in green. Glycan at N343 represents by dark cyan ball and stack.

Antibodies have a major role in the immunological response to SARS-CoV-2 infection. RBD-specific or N-terminal domain (NTD)-specific mAbs of S-protein have been shown to be the most promising and effective antiviral therapeutics for COVID-19.^11,12^ However, mAbs still confront significant challenges, with the most pressing concern being whether mutations of RBD Omicron subvariants can confer antibody resistance. Particularly, RBD is immunodominant and contains WT epitopes for 90% of the antibodies elicited by natural infection or vaccination.^13^ Thus, the distributions of RBD Omicron mutations on these epitopes could potentially disrupt biological characteristics of RBD epitope surfaces, leading to escape from the majority of mAb therapeutics and/or compromise the acquired human immune profile. While this is a hot topic with valuable studies using different approaches,^14–23^ further research effort is necessary to probe the consequences of RBD Omicron mutations on antigenic profiles, particularly those that address the molecular dynamics and thermodynamic aspects, which is the goal of the current study.

RBD-targeting mAbs are typically classified into four classes (class 1 to 4, which we utilized here)^24^ or six RBD receptor binding sites (RBSs)^25^ depending on various criteria such as their binding modes to “up”, “down”, or both RBD conformations, competitions with angiotensin converting enzyme-2 (ACE2) receptor, the location of their RBD epitope, and derivation from antibody gene families. These two classifications are highly overlapping. Class 1 mAbs are mainly derived from the VH3-53 or VH3-66 germlines with short CDRH3 (complementarity-determining region 3 of heavy chain) loops that block ACE2 and exclusively bind to “up” RDB conformation.^24^ Class 2 mAbs are encoded by different germ segments and has a long CDRH3 loop and directly compete with ACE2 by binding to both “open” and “close” RBDs. Unlike classes 1 and 2, classes 3 and 4 bind the RBD but do not interfere with ACE2-binding site. Class 3 recognizes both RBD conformations, whereas class 4 binds only the “up” RBD.

In this study, the molecular basis of antibody binding against emerging Omicron subvariants such as BA.1, BA.2, and BA.4/5 is investigated and compared with its WT to determine the overall impact of RBD mutations or the effect of each specific RBD mutation. To achieve this, the binding of ten mAbs against RBD WT or Omicron subvariants (BA.1, BA.2, and BA.4/5) has been probed. These mAbs mainly recognize and target the RBD class 1 (BRII-196^26^, CT-P59^27^, and LY-CoV016^28^), class 2 (LY-CoV555^29^, COV2-2196^30^, and REGN10933^31^), class 3 (REGN10987^31^, COV2-2130^30^, and S309^32^) and class 4 (S2X259^33^) epitope regions (**Figure 2**). These mAbs cover almost all epitope sites on the RBD. In total, 40 different mAb-RBD complexes, i.e.,10 mAb for each RBD case, have been investigated using molecular dynamics (MD) simulations and followed by end-point binding free energy (BFE) calculation with molecular mechanics Generalized Born surface area (MM-GBSA) method. Such investigations are crucial not only for understanding the molecular and amino acid origins of immune evasion by these Omicron subvariants but also for instructing better design of mAbs and vaccines.

**Figure 2.**
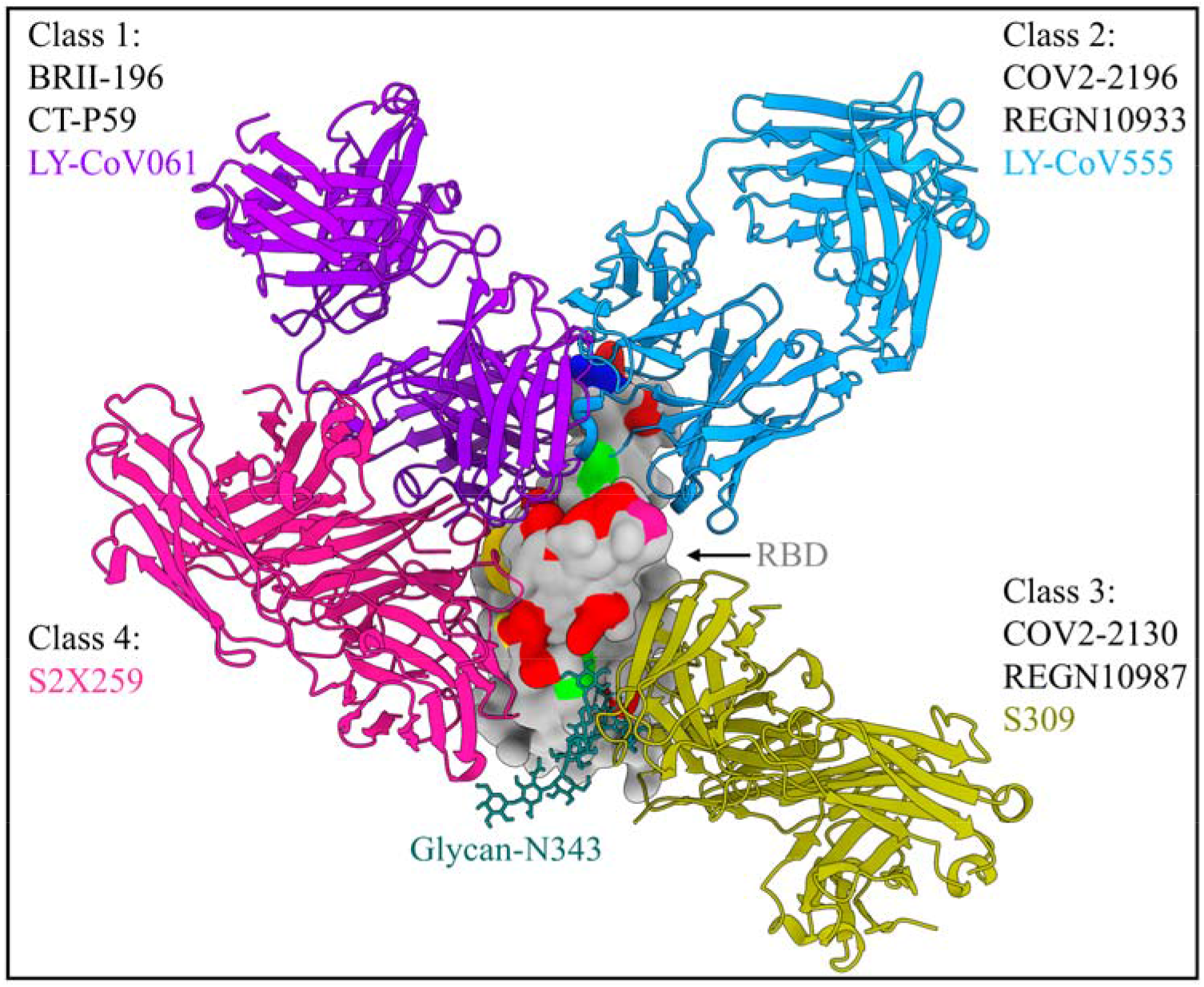
Schematic illustration of different RBD-binding epitopes targeted by four distinct classes of RBD-specific mAbs, along with specific examples of the interface bound Fab mAb-RBD complex in each class. The representation of RBD (gray surface) with the Omicron mutations is the same as in Figure 1. The Fab mAb is represented by ribbon.

## 2. Results and Discussion

To map escape mutations and determine the effects of RBD Omicron subvariant mutations as well as to elucidate the molecular basis for immune recognition of RBD antigenic sites, we explicitly generated four different RBD-Fab models for each individual mAb. The first model is for WT RBD with the fragment antigen-binding (Fab) region of the mAb, while the remaining three models are for BA.1, BA.2, and BA.4/5 RBD with the Fab. The comparison of the Omicron models against the WT model allows us to predict the mutational implications of each Omicron subvariant and how Omicron has evolved to dodge acquired immunity. In total, this study targeted forty mAb-RBD complexes, i.e., ten mAb × four different RBDs.

We developed a procedure for creating initial structural models having all RBD Omicron mutations as fully described in Supporting Information (SI). **Table S1** summarized all 40 solvated models of the glycosylated mAb-RBD complex in WT or Omicron variants. The MD and MM-GBSA protocols are also described in SI.

### 2.1 Conformational changes in RBD Omicron subvariants impair the stability of the mAb-RBD complex

To gain insight into how the Omicron mutations impact the dynamic behavior of mAb-RBD complex, two replicates of all-atom MD simulations were run over 200 ns for forty biosystems and their MD trajectories are monitored to verify the structure stability of Omicron subvariant complexes vs related WT. **Figure 3 (a)** (top panel) displays the averaged root-mean-square-deviation (RMSD) of the complex from these two MD runs, while **Figure S1** shows the time evolution of RMSD for all models. These figures, with a few exceptions, show that the mAb forms a fairly unstable complex with Omicron subvariants, as evidenced by relatively large fluctuations in RMSD with many plateaus relative to the WT.

**Figure 3.**
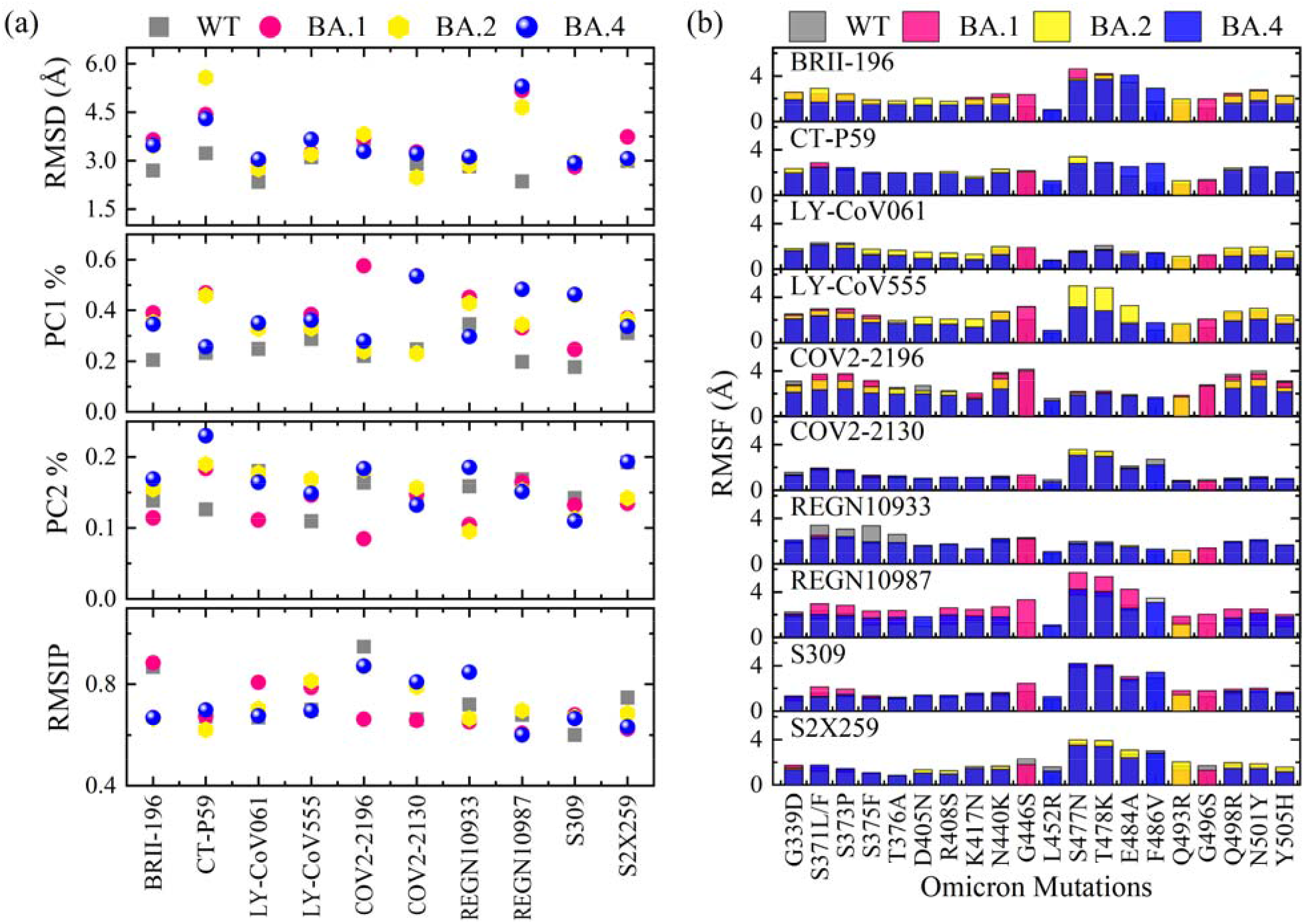
The dynamic profiles and conformational changes in the RBD Omicron subvariant complexes with mAb. The top panel of (a) shows the averaged root mean square deviation (RMSD) of the heavy atoms of mAb-RBD complexes obtained from two independent MD runs. For better visual clarity, the standard deviation of error (SD) from two runs is not shown. Except in CT-P59_BA.2 or BA.4/5_, SD does not exceed 0.5 Å in all systems. Middle panels of (a) represent the contribution of the lowest PC1 and PC2 modes to the total conformational fluctuations for only RBD residues (T333-P527) and how the Omicron mutations explore a larger conformational space than in its WT. Bottom panel of (a) illustrates the MD reproducibility from two replica simulations using the root mean squared inner product (RMSIP) for the PC1 and PC2 of whole heavy atoms of the mAb-RBD complex. (b) Local conformational and stability of RBD Omicron mutations with respect to their WT based on the root-mean-square fluctuations (RMSF). SD does not exceed 0.8 Å. All analyses are based on 5000 snapshots taken from each MD run over the course of 200 ns.

Additionally, CT-P59 and REGN10987 have relatively large RMSD compared to other complexes, especially when their RBD contains Omicron mutations. This result implies that RBD Omicron mutations may change the overall stability of the mAb-RBD complex. To demonstrate that, the principal component analysis (PCA) has been performed to extract the fundamental motions of RBD in Omicron subvariants vs WT. The first two principal components (PC1 and PC2) of only RBD (T333-P527) in different Omicron seniors is projected onto their WT RBD as shown in **Figure S2**. PCA shows notable variation in collective motion for Omicron RBDs comparing to WT, indicating conformational changes between them. The PC1 of Omicron RBDs captures relatively largest variance during the MD simulations comparing to WT, clearly support our finding that Omicron RBDs explores a relatively larger conformational space than in WT (see the amount (%) of variance accounted by PC1 and PC2 in the middle panels of **Figure 3(a)**). Analysis of the root-mean-square-fluctuations (RMSF) further confirms this observation (**Figure 3(b)** and **Figure S3**). In most cases, the RBD Omicron subvariants exhibit an increase in overall flexibility, particularly in the β5–β6 loop region (residues 473-488) of RBM. It is possible that the high flexibility of these Omicron sites and the mAb residues at the interface resulted in a destabilized complex, which in turn reduce the binding of the mAb-RBD Omicron complexes in comparison to their corresponding WT system. Collectively, our findings show that mAb-RBD Omicron complexes are relatively less stable and more dynamic than their WT counterparts. As a result, immune evasion of Omicron subvariants could be facilitated, limiting mAb efficacy.

Although the simulation time is only 200 ns, there is a considerable overlap between two MD runs as seen by **Figure S1** of the RMSD. This is a good indication that the independent MD runs are reproducible. This is even more pronounced by both the PCA for whole complex (**Figure S4** for mAbs class 1 as an example) and the value of the root mean squared inner product (RMSIP) for the PC1 and PC2 that ranges from 0.65 to 0.9 (**Figure 3(a)** bottom panel).

### 2.2 RBD Omicron subvariant mutations reduce or abolish mAb binding by changing their electrostatic potential surface

BFE is a critical thermodynamic quantity that drives all molecular processes, including the mAb-RBD binding mechanism. It measures the strength of the interaction between the mAb and its antigen like RBD and is thus often directly related to the mAb’s potency. In present study, MM-GBSA approach was used to predict the complete BFE profile between the mAb and RBD at 310 K (37 °C) under a neutral pH and 0.15 M univalent NaCl salt concentration. We first compare our predicted BFE of the WT mAb-RBD complexes to the available experimental results,^26–29,32,34–36^ as listed in **Table S2. Table S2** shows that the calculated BFE using MM-GBSA has excellent agreement in six complexes, but it is underestimated in BRII-196_WT_ and overestimated in LY-CoV016_WT_, S309_WT_, and S2X259_WT_. Overall, there is an acceptable correlation between the MM-GBSA and experimental values (the Pearson correlation r = 0.63 for all 10 models as shown in **Figure S5**). When the computed BFE values for BRII-196_WT_, LY-CoV016_WT_, S309_WT_, and S2X259_WT_ are excluded, the correlation coefficient increases to 0.93 (**Figure S5**). This observation is not surprising, since the MM-GBSA method is highly dependent on the system under investigation and is influenced by many parameters such as the solute dielectric constant, MD length and replica, the ensemble used to calculate entropic contributions, and so on.^37^ The experimental values, on the other hand, fall within a relatively narrow range of -10.5 to -17 kcal/mol, making the linear fit with this computational method is significantly affected when there is a little drift. Nonetheless, it is well known that the MM-GBSA can provide a good BFE ranking that agrees reasonably well with the experimental binding affinity ranking.^38^ Additionally, this popular MM-GBSA methodology offers a tradeoff between accuracy and efficiency.^37,39,40^

To assess the effects of Omicron mutations on mAb binding, the BFE between different Omicron RBDs and mAb is calculated and compared to the binding of related WT complex, as shown in **Figure 4**. Roughly speaking, a positive BFE indicates that there is no spontaneous association between the mAb and RBD (not favorable or no binding), whereas a negative value reflects the association (favorable). **Figure 4** shows that the BFEs for BRII-196, CT-P59, COV2-2196, and LY-CoV555 are completely flipped toward positive values in all Omicron cases, suggesting that they have all lost their ability to neutralize RBD Omicron subvariants. This finding is in consistent with recent experimental observations.^14,15,41^ REGN10987 also loses its ability to associate with BA.1 while retaining moderate binding to BA.2 and BA.4/5. A recent experimental studies support this observation.^19,41^ The bindings of REGN10933 and LY-CoV016 to BA.1, BA.2, and BA.4/5 are substantially dropped, with the reduction in BFE ranging from 30 to 68% of their WT complexes. On the other hand, the COV2-2130 exhibits a moderate reduction in BFE to Omicron subvariants especially of BA.1, ranging from 10 to 5 % compared to its WT. This again also in line with previous study.^14,19^

**Figure 4.**
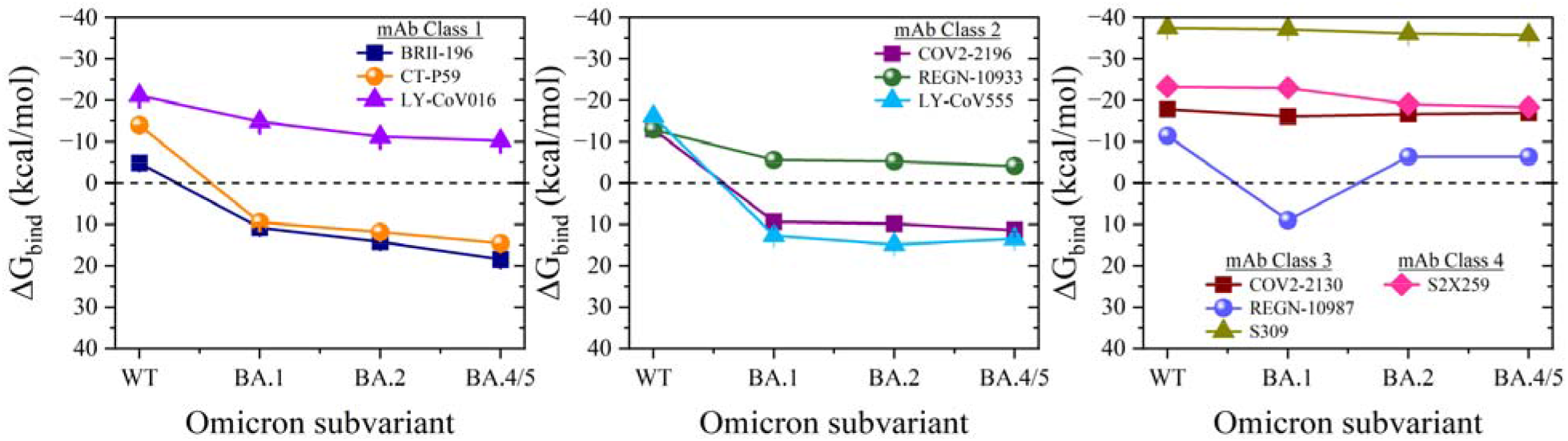
Impact of RBD Omicron mutations on the binding of broad mAb in all four classes. The standard error of mean (SEM) along the 200 ns from both runs are less than ± 0.5 kcal/mol.

S309 and S2X259 preserve their ability to bind BA.1 but are reduced against BA.2 and BA.4/5, particularly S2X259. The drop in binding for S2X259 against BA.2 and BA.4/5 is 18 and 21% of their original WT, respectively. The binding behavior of the S309 and S2X259 against Omicron subvariants is good correlated with experimental data. ^14,41,42^

In general, **Figure 4** shows that the binding of mAb classes 3 and 4 are less affected by Omicron than mAb classes 1 and 2.

To gain a better understanding of the primary driving force that causes Omicron subvariants to weaken/lose mAb binding, the BFE is dissected into its energetic components as shown in histogram plots of **Figure 5** for 9 mAbs as well as **Figure S6** for S2X259. Then, the relative energetic components with respect to relevant WT system is estimated as shown in doughnut plots of **Figure 5** and **Figure S6**. These figures indicate that the source of escape Omicron mutations from a broad mAbs mainly originates from changing their electrostatic interactions. Obviously, seven RBD Omicron mutations, G339D, N440K, L452R, T478K, Q493R, Q498R, and Y505H, change from polar or nonpolar residues to charged residues, whereas K417N, D405N, R408S, and E484A have opposite charge modifications (change from charged residues to polar/nonpolar). These charge modifications undoubtedly have induced the electrostatic changes between Omicron RBDs and mAbs as demonstrated by quantitative and qualitative results in our study. Consequently, the recognition and specificity of mAb are strongly affected by these electrostatic changes. This observation is fully consistent with other studies.^3,43,44^

**Figure 5.**
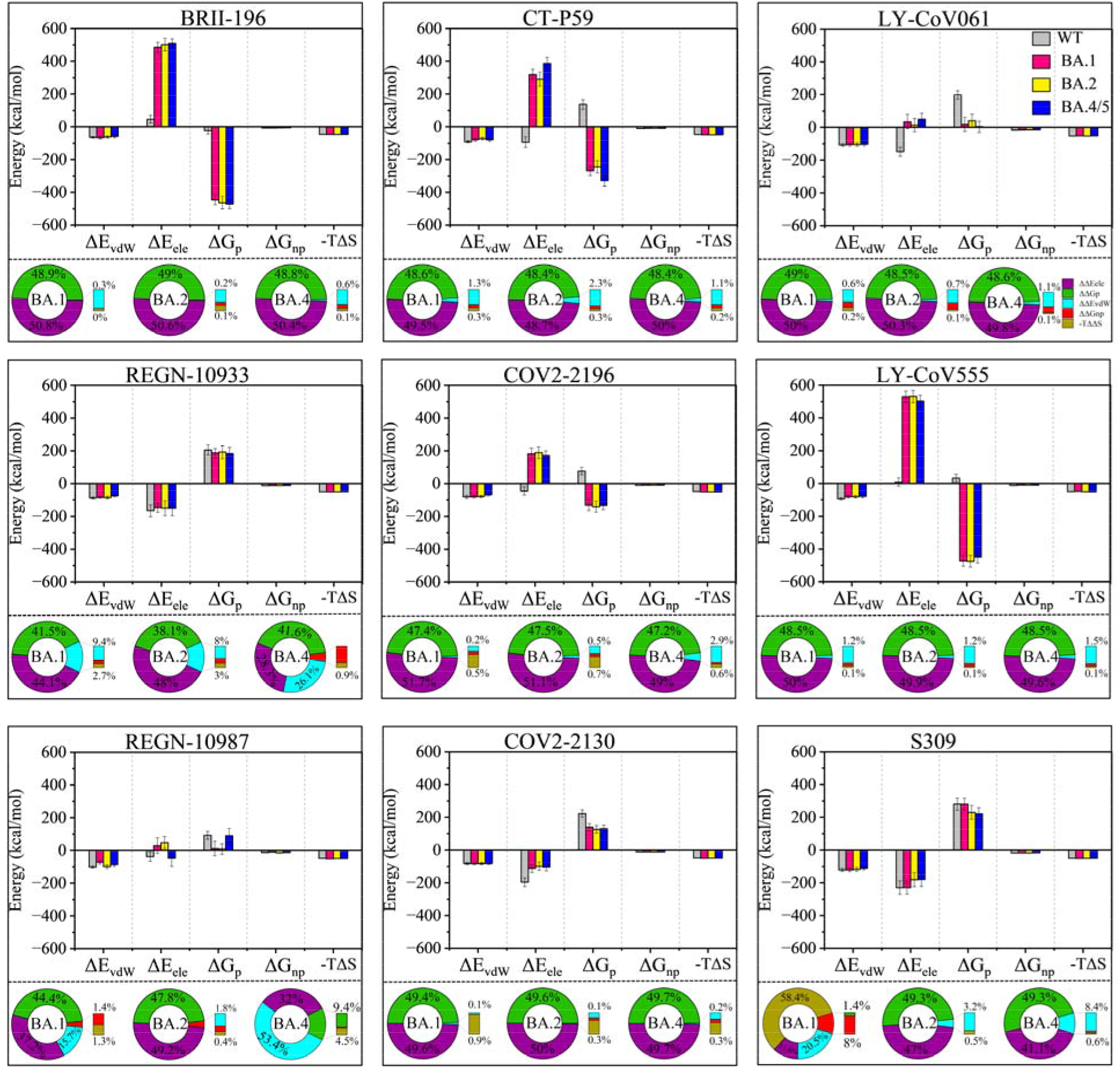
The electrostatic interaction role of Omicron subvariant mutations in conferring mAb resistant. For each panel, the histogram displays the BFE dissection in terms of energetic components including van der Waals (ΔE_vdW_), Coulombic electrostatic (ΔE_ele_), electrostatic polar (ΔG_p_), non-polar (ΔG_np_), and entropic (-TΔS) contributions. Because the standard error of mean (SEM) is too small to appear in this histogram, the standard deviation (SD) is reported instead. The doughnut plots in the bottom of each panel represent the relative energetic differences of Omicron subvariants with respect to its WT value.

Along with these, changes in van der Waals (ΔE_vdW_) interaction also contribute to the loss/reduction of binding of specific mAbs such as CT-P59, LY-CoV061 and LY-CoV555 in all Omicron subvariants, as well as S309 binding in BA.2 and BA.4/5. In BA.4/5, although vdW interaction changes are not large as electrostatic components, they are significantly implicated in the BRII-196_BA.4/5_, COV2-2196_BA.4/5_ and REGN10933_BA.4/5_ loss/decrease binding. This indicates that either L452R or F486V could impair the hydrophobic contacts or steric hindrances for these complexes (see section 2.3).

### 2.3 Consequences of Omicron mutations extend beyond exact substitutions to influence the interactions of interface mAb residues and local and non-local conserved RBD residues

The subtle impacts of each RBD Omicron subvariant mutations on mAb were distinguished by using the per-residue BFE decomposition. With this analysis, the energy contribution of a single residue can be computed by summing its interactions across all residues in the complex.

First, the WT RBD epitopes for each mAb are identified, but this time based on their thermodynamic energies, and compared with RBD epitope from the static structural studies using the IEDB database^45^ as shown in **Table S3**. Remarkably, the energetic analysis at the residue level predicts not only the previously identified RBD epitope, but also additional residues for each mAb (color sites in **Table S3**). This table also revealed a high number of Omicron mutations at certain mAb binding sites, which are as following: BRII-196_WT_: 3 (5); CT-P59_WT_: 6 (6); LY-CoV016_WT_: 7 (8); LY-CoV555_WT_: 3 (4); COV2-2196_WT_: 5 (7); COV2-2130_WT_: 5 (5); REGN10933_WT_: 4 (8); REGN10987_WT_: 3 (3); S309_WT_: 1 (2); and S2X259_WT_: 6 (7) mutations based on structure or (current) studies (see bold sites in **Table S3**). **Figure 6** display the relative per-residue BFE for each Omicron mutation (ΔΔG_per-residue_) that is estimated as the energy difference between a mutated site and its WT. It is expressed as ΔΔ*G*_per–residue_ = (ΔΔ*G*_per–residue_)_*BA*.*i*_ – (ΔΔ*G*_per–residue_)_*WT*_, where *BA*.*i* can be BA.1, BA.2 or BA.4/5.

**Figure 6.**
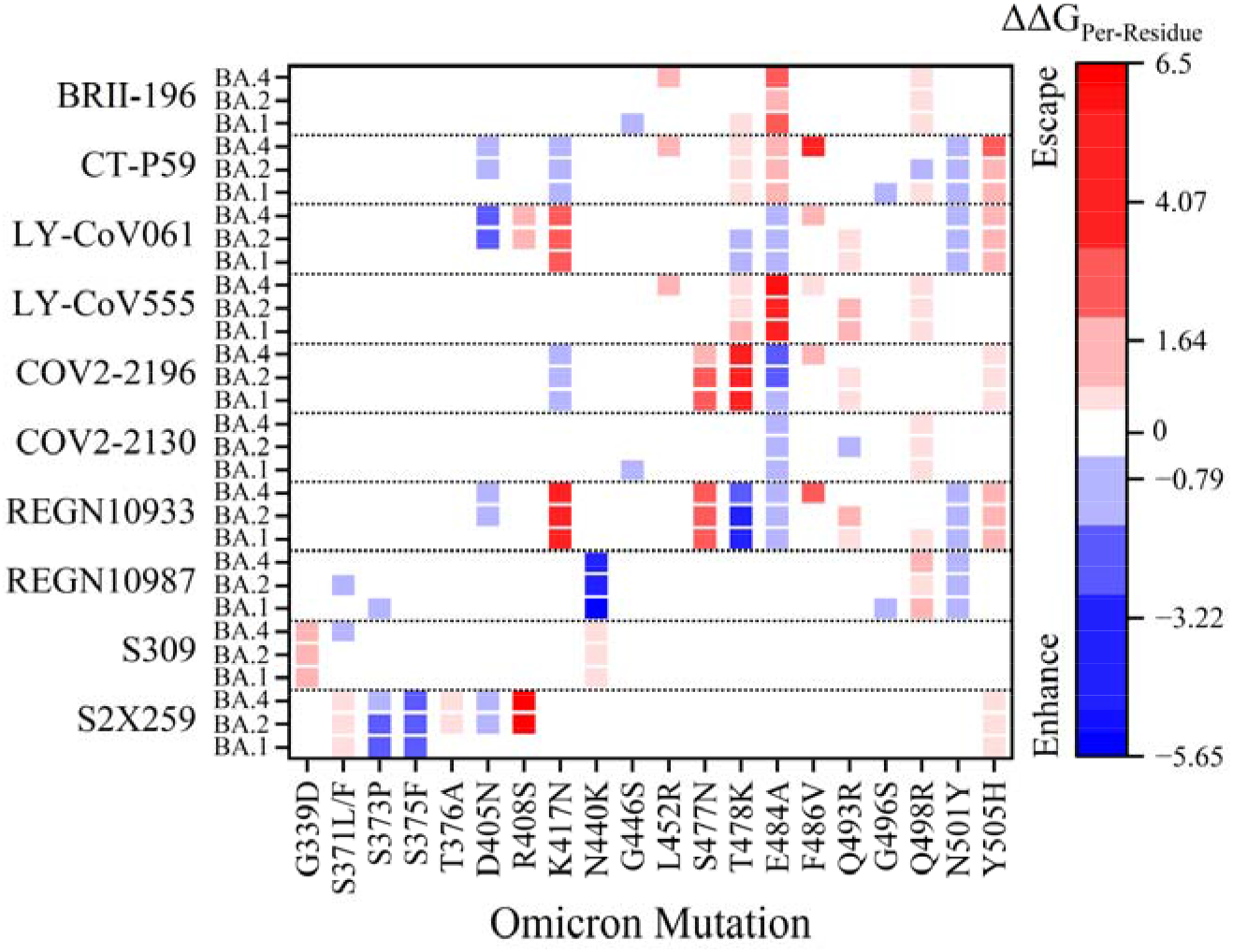
Molecular interaction footprint map of Omicron mutations on different mAb classes. These analyses are based on the relative per-residue BFE (ΔΔG_per-residue_ in kcal/mol unit) of each Omicron mutation with respect to its WT. Mutations that escape antibody binding are represented by more intense red cells, while those that enhance binding or have no binding effect are represented by blue or white cells, respectively.

Some Omicron mutations can be identified as escape mutations including K417N from LY-CoV061 and COV2-2130, S477N from COV2-2196 and REGN10933, and importantly E484A from BRII-196, CT-P59, and LY-CoV555. Interestingly in BA.2 and BA.4/5, LY-CoV061 and S2X259 share the same escape mutation R408S. In BA.4/5, L452R confer the immune resistance against BRII-196, CT-P59 and LY-CoV555 as well as F486V in CT-P59, LY-CoV061, COV2-2196, and REGN10933. Other mutations could be classified as potential escape mutations such as Q493R, Q498R and Y505H. However, not all RBD Omicron mutations facilitate immune evasion. For instance, S373P and S375F in the S2X259 complex or T487K in the REGN10933 enhance the binding. This is also true for N501Y for some mAbs. Other mutations, such as E484A against COV2-2196 or D405N against S2X259, appear to enhance binding, but the reality is quite the opposite, as we will see in next section. Therefore, the per-residue BFE decomposition of only the actual mutation site is inadequate to describe the complete picture of mutational impacts. It requires the incorporation of unmutated RBD residues whose biological environment has changed due to mutations, particularly those adjacent to the mutation, as well as mAb residues that form a pair with WT RBD as shown in **Figure S7. Figure S7** shows all possible per-residue BFE differences of all unmutated RBD or mAb residues that are above or less than ±0.15 kcal/mol. **Table 1** summarized the key residues in RBD or mAb chains that are significantly affected by Omicron mutations with range differences of more or less than ±1 kcal/mol.

**Table 1.**
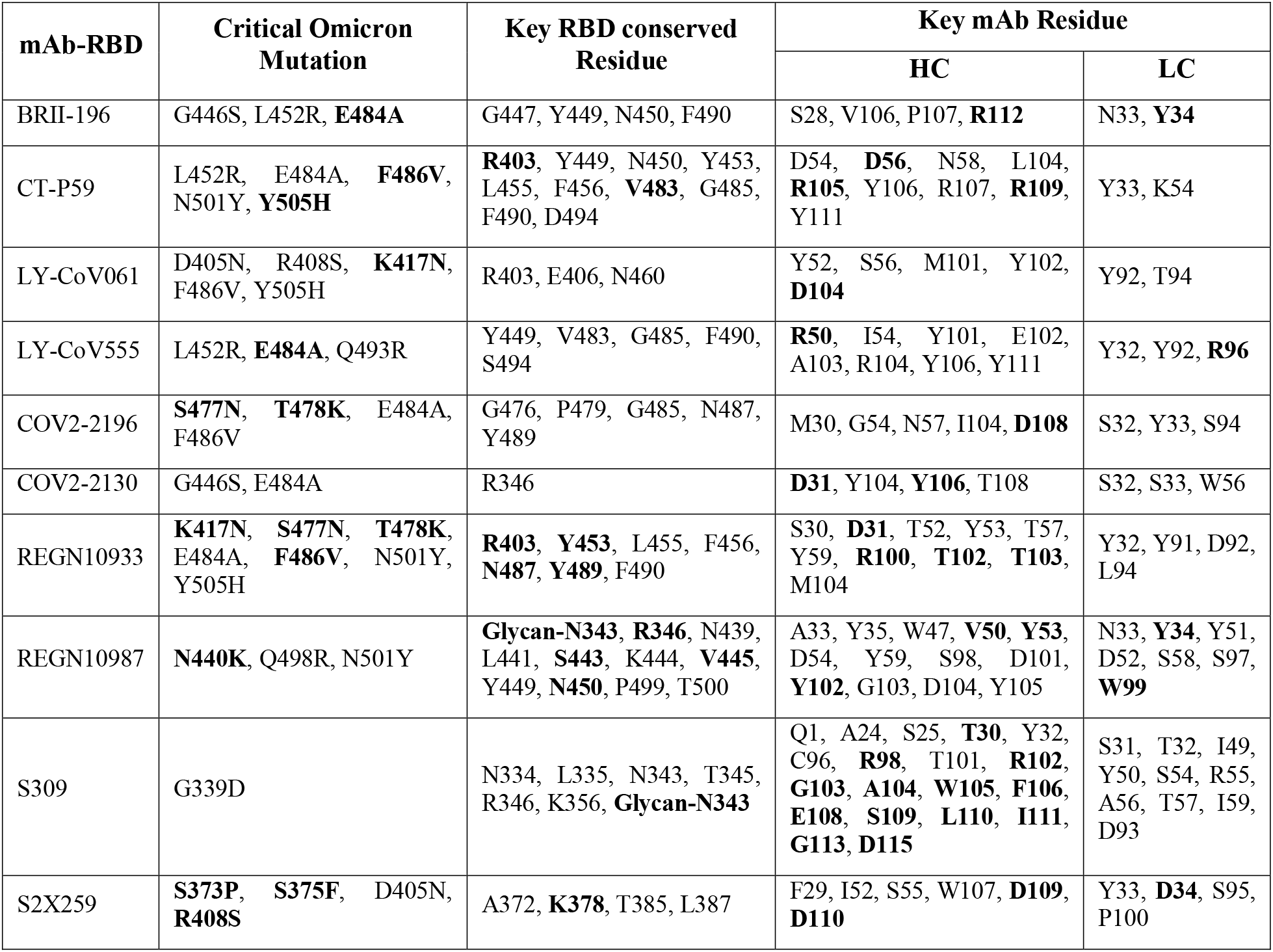
A summary of the major Omicron mutations that affect mAb binding through impact interaction contribution due to substituting at the actual mutation site, local and non-local impact on RBD conserved residues, or the intermolecular interaction impacts on the heavy and light chain (HC or LC) sites of mAb. This identification is based on per-residue BFE differences that are greater than 1 kcal/mol (drop binding) or less than -1 kcal/mol (enhanced binding). The most significant differences with ≥ 2.5 kcal/mol or ≤ -2.5 kcal/mol are labeled in bold.

Obviously, the effect of RBD Omicron mutations extends to more than actual substations at a certain site. In general, many other conserved RBD or mAb chain residues are implicated in the consequences of evading each mAb. HC residues of mAb are more involved in these Omicron ramifications. Significantly, the per-residue BFE differences for some of these key residues are very critical and reached more or less than ±2.5 kcal/mol (see bold residues in **Table 1**). These critical residues could be mainly contributed to conferring antibody evasion (see next section). Furthermore, the interacting spectrum for the mAb that its binding is less affected by Omicron mutations such as COV2-2130 differs from that of its WT. This indicates that while the net BE of some mAbs remains significantly unchanged, their intermolecular interactions with Omicron are not completely identical. It should also mention that the glycan at N343 is anticipated in the interactions with mAb class 3, in particular S309 in all cases and REGN10987_BA.2_ and REGN10987_BA.4/5_. In the S309 complexes, the impact of Omicron mutations on their interactions with glycan did not follow a specific trend, in which BA.1 mutations can enhance interactions with glycan while BA.2 or BA.4/5 mutations can do the opposite. In REGN10987, both BA.2 and BA.4/5 are made many interactions with glycan that are not observed in both WT and BA.1. Further investigations are required here. The data in **Table 1** does not mean that other residues in **Figure S7** with less significant BFE differences can be ignored.

At this stage, one can infer that the impacts of Omicron mutations on mAb binding are not limited to just the impact on the actual interactions of the mutated site alone but extend beyond the local and non-local unmutated RBD residues and the mAb residues involved in the interface interactions. More details about the structural and energetic consequences of the Omicron mutations will be discussed in next section.

### 2.4 Mapping AA-AA pair interactions and Structural basis of the mAb-RBD Omicron

In-depth investigations were first carried out pairwise BFE decomposition to quantitatively capture the actual interacting AAs of mAb-RBD complex and to determine the molecular basis of reduced/lost binding of mAb to Omicron RBDs. In each mAb, we first accounted all potential pairs in all Omicron cases and WT, and then we calculated the change in the pairwise BFE decomposition for each pair with respect to its WT pair as shown in **Table S4**. Here, differences larger than -0.15 or less than 0.15 kcal/mol are ignored. The pairwise BFE decompositions were backed by intermolecular interaction analyses such as hydrogen bonding (HB), salt-bridge, and others to reveal their structural basis. The changes in HB profiles in Omicron complexes compared to WT are fully included in the **SI excel sheet**.

Omicron mutations at the mAb-RBD interface result in steric clashes and/or the abolition of salt-bridge and/or HBs, as summarized below for each mAb. Here, mAb residues are distinguished by the subscripts H or L, which denote the heavy or light chains of mAb, respectively.

#### BRII-196

Our findings indicate that the E484A mutation is a major source of evading binding of BRII-196 against Omicron subvariants due to the loss of salt bridges with R112_H_ (**Figure 7**) and weak strength of the pairs with N33_L_, Y34_L_, V106_H_, P107_H_ from CDRL1 and CDRH3 respectively. In contrast, E484A slightly enhances the interactions with E52_L_ and K55_L_ from CDRL2. Besides, it has either a positive or negative effect on the interactions of the vicinity conserved residues, particularly those of V483 and F490 (**Table S4**). In BA.1, G446S can form more stable HBs with Y27_H_, S28_H_, and S31_H_. However, this mutation impairs the interactions of nearby conserved residues, particularly those of V445 and G447, and causes steric clashes with BRII-196. In BA.4/5, L452R could disrupt the hydrophobic interactions between Y449, L452, and F490 from RBD and CDRH1/H3 residues of BRII-196 (**Table S4**), avoiding the binding even more.

**Figure 7.**
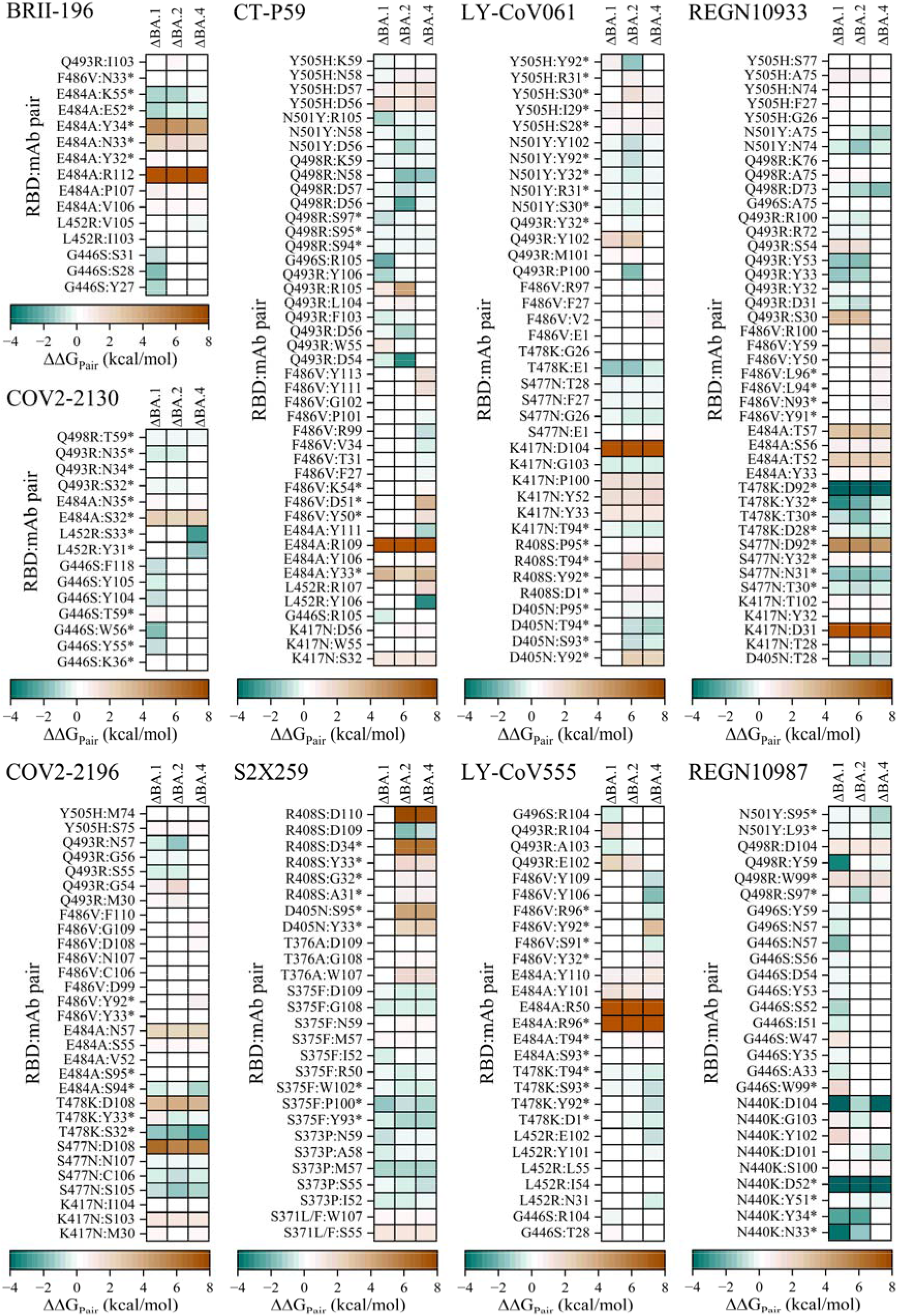
The impact of the RBD Omicron mutations on the AA-AA pair interactions between RBD and mAbs. This analysis is based on the differences (ΔΔ*G*_pairwise_ = (ΔΔ*G*_pairwise_)_*BA*.*i*_ – (ΔΔ*G*_pairwise_)_*WT*_) in the Pairwise BFE decomposition for each pair of the Omicron RBD-mAb and its corresponding WT complex. ΔBA.1, ΔBA.2, or ΔBA.4 represent the change in the pairwise BFE with respect to WT. This figure shows only the actual AA-AA interaction differences of the mutated site that forms pair with certain mAb and has energetic differences energy larger than 0.15 or less than -0.15 kcal/mol. The mAb light chair residues are labeled with asterisk to distinguish from heavy chain residues.

#### CT-P59

Since CT-P59 has a different binding mode and unique RBD epitopes than other class 1 mAbs,^27^ E484A is also a dominant mutation that loses the strong salt-bridges with R109_H_ and anion-π interaction with aromatic rings of Y33_L_ (**Figure 7**). K417N and Y505H prevent binding even more by excluding HBs and possible hydrophobic contacts with S32_H_ and D56_H_, respectively. While Y505H also weakens interactions with D57_H_ and N58_H_, particularly in BA.2 and BA.4/5, N501Y or Q498R could balance this loss by forming stronger pairs with these CRDH2 residues (**Figure 7**). In BA.1, G496S strengthens the pair with R105_H_. This also enhances contacts between the conserved residue Y495 and R105_H_ but has a negative impact on the Y449 and R105_H_ pair (**Table S4**). In BA.1 and BA.2, Q493R abolishes the HB with R105_H_ and weakens pairs with W55_H_ and L104_H_ but it leads to strong interactions with D54_H_, D56_H_ and Y106_H_. However, it has an impact on neighboring S494 interactions including its HBs. Although the D405N mutation of BA.2 and BA.4/5 is not engaged in the interaction with CT-P59, it may destabilize the R403, thereby impeding the salt bridges with D56_H_. In BA.4/5, even though L452R can form the π-cation and/or hydrophobic interactions with the aromatic sidechain of Y106_H_, it has two adverse consequences. First, it electrostatically repels with R107_H_. Second, it negatively influences the hydrophobic interactions involving the unmutated residues F490 and L492 with Y106_H_. Unlike the midst impact of L452R, the F486V mutation is responsible for further hindering BA.4/5 from binding to CT-P59, which it fails to form the HBs with D51_L_ and Y111_H_ and/or the hydrophobic contacts with Y50_L_ and Y113_H_.

#### LY-CoV061

K417N is mainly responsible for reducing binding with LY-CoV061 in all Omicron subvariants by losing the salt-bridge with D104_H_ as well as disrupting the interactions with Y33_H_, Y52_H_ and P100_H_ of its heavy chain (**Figure 7**). S477N maintains the same interactions with E1_H_, G26_H_, F27_H_ and T28_H_ with slight strength differences. T478K strengthens the interaction with E1_H_ while weakening it with G26_H_. Y505H has fewer packing interactions with S28_L_, I29_L_, S30_L_, and R31_L_ light chain residues, whereas N501Y compensates for some of them (**Figure 7**).

In BA.1 and BA.2, Q493R more impairs LY-CoV061 binding either by steric overlaps with CDRH3 or inability to form HBs with its Y102_H_ (**Figure 7** and **SI excel sheet**). In BA.2 and BA.4/5, R408S additionally contribute to attenuate binding with its light chain by breaking the HBs with T94_L_ and diminishing the interactions with D1_L_, Y92_L_, and P95_L_ (**Figure 7**). Therefore, R408S could be an escape mutation for LY-CoV061. Despite the lack of HBs with Y92_L_, D405N in BA.2 and BA.4/5 might still maintain the interactions with S93_L_, T94_L_ and P95_L_ from CDRL3. In BA.4/5, L452R has no binding impact, but F486V renders a further drop in binding owing to lowered interactions with E1_H_, V2_H_, F27_H_, as well as an unfavorable interaction with R97_H_ (**Figure 7**).

#### LY-CoV555

In all Omicron subvariants, E484A is the main source of complete loss of LY-CoV555 binding due to collapse the salt-bridges and HBs with R50_H_ and R96_L_, as well as the HBs with Y101_H_ and Y110_H_ (**Figure 7** and **SI excel sheet**). It also influences the interactions of adjacent conserved residues with LY-CoV555, particularly V483 (**Table S4**). Overall, the change in surface charge distribution of E484A from highly negative charge to polar AA causes a significant disruption in electrostatic interaction not only at the actual 484 site but also at its nearest neighboring sites. Here, *ab initio* quantum methodologies could provide a more accurate description of its partial charge distributions.^46–53^ T478K is still paired with Y92_L_ and S93_L_, with no discernible effect.

In BA.1 and BA.2, Q493R destabilizes its binding by abolishing HB with R104_H_, in addition to adversely impacting the strength of pair with E102_H_. The HB network between S494 and R104_H_ is also disrupting by this mutation (**SI excel sheet**). For BA.4/5, our findings revealed that L452R could also interact with I54_H_ and L55_H_, but at a lower strength due to possible steric clashes. L452R can also make more favorable interactions with N31_H_, Y101_H_ and E102_H_, but it could form many unfavorable long-range electrostatic interactions with positively charged LY-CoV555 residues in the surrounding environment. Besides that, it required a large amount of desolvation energy to stabilize its interface with this mAb. Therefore, the L452R net difference is energetically unfavorable, which is consistent with previous studies.^54,55^ F486V, on the other hand, abrogates the strong HB with Y92_L_ and reduces hydrophobic interaction with Y31_L_, but it is slightly tighter with S91_L_, R96_L_, Y106_H_, and Y109_H_.

#### COV2-2196

Even though S477N and K478K could relatively form stronger pairs with S105_H_ and S32_L_ respectively, they are the two major Omicron mutations involved in the total loss of binding to COV2-2196 due to breaking the O…OH HBs with D108_H_ (**Figure 7**). Furthermore, these mutations disrupt the interactions of the conserved residue G476 with D108_H_ and P479 with S32_L_ and Y33_L_, as well as the HBs between N487 and D108_H_ (**Table S4** and **SI excel sheet**). Besides that, the loss of the pair with S55_H_ and the absence of HBs with N57_H_ suggest that E484A may hinder its binding further. This impact, however, may partially compensate by generating the HB between A484 and S95_L_ and its favoring interaction with S94_L_ (**Figure 7**). K417N slightly weakens the pairs with M30_H_, S103_H_, and I104_H_. Y505H has a marginal effect because it misses the weak pairs with the M74_H_ and S75_H_. In BA.1 and BA.2, Q493R also reduces the binding due to not participating in the HB with G54_H_. While the F486V of BA.4/5 significantly damps the interactions with Y92_L_, D99_H_, and N107_H_ to F110_H_ (**Figure 7**).

#### COV2-2130

E484A eliminates the HB with S32_L_ and fails to pair with N35_L_, resulting in lower COV2-2130 binding. N440K retains the same pairs with G110_H_ and P111_H_. In BA.1, G446S could relatively strengthen the pair with Y55_L_, W56_L_, Y104_H_, Y105_H_, and F118_H_ but weaken with T59_L_ (**Figure 7**) and could destabilize the interaction of the adjacent conserved residues V445 and Y449. Since Q493R preserves the same pairs with the CRDL2 loop (residues S32_L_ to N35_L_) in BA.1 and BA.2, it has no major effect. In BA.4/5, L452R can form HBs with Y31_L_ and S33_L_, enhancing the overall interactions with COV2-2130.

Since the conformation of the conserved residue R346 differs between the WT and Omicron subvariants, its interactions are expected to be distinct. Indeed, our findings reveal that the R346 in WT RBD makes more favored interactions with D31_H_, S53_H_, and I55_H_ than R346 in Omicron RBDs (**Table S4**). In contrast, R346 in Omicron RBDs interacts strongly with W33_H_, Y106_H_, D107_H_, T108_H_. Overall, R346 could reduce the COV2-2130 binding in Omicron subvariants. However, the favored interaction of T345 with T108_H_ in Omicron subvaraints may partially offset such binding reduction.

Interestingly, the extremely stable networks of the critical K444 conserved site with CRDH3 residues are still intact (**SI excel sheet**), which could be the main reason why Omicron subvariants have little effect on COV2-2130 binding.

#### REGN10933

K417N loses charge interactions with D31_H_, which mainly contributes to the decreasing affinity of REGN10933 (**Figure 7**). This increases the chances that the conserved residue R403 of the Omicron RBDs can pair with D31_H_ and partially compensate for such loss, particularly in BA.2 and BA.4/5, where D405N is suspected to play a role (**Table S4**). However, this possible pair reduces the interactions between Y453 and D31_H_ and S30_H_. S477N eliminates HBs with D92_L_, but this loss is totally overcome by the formation of an ionic pair of T478K with D92_L_ in the presence of HBs (**Figure 7**). In addition, S477N can slightly reduce/increase interaction with T30_L_, N31_L_, and Y32_L_ while T478K can form a HB with Y32_L_ and relatively more favor interactions with D28_L_ and T30_L_. As a result, S477N has an adverse effect on the REGN10933, whereas T478K has a positive effect. E484A also has a detrimental effect on the binding because it ignores interactions with Y33_H_, T52_H_, Y53_H_, S56_H_, and T57_H_. Likewise, Q493R and Y505H can impair binding, whereas Q498R has no overall effect because it can counterbalance the lose interactions with A75_H_ and K76_H_ by gaining interaction with D73_H_. N501Y could little enhance its binding especially with N74_H_, and A75_H_. In BA.4/5, F486V can diminish the hydrophobic interactions with the light and heavy chain residues including Y91_L_, N93_L_, L94_L_, L96_L_, Y50_H_, Y59_H_, and R100_H_. Even without the mutation at this position, the HB between F486V and Y59_H_ is lost, suggesting that E484A mutation may have an impact on this pair rather than the actual F486V mutation. Altogether, the REGN10933 binding against BA.4/5 can decline even further.

Omicron mutations may also affect the interactions of the conserved residues Y421, L455, F456, A475, G476, G485, N487, and Y489 with REGN10933.

#### REGN10987

In all Omicron cases, N440K strengthens the binding by forming salt bridges with D52_L_ and ionic interactions with D104_H_, but it weakens with S100_H_, Y102_H_, and G103_H_ (**Figure 7**). Further, this mutation results in abolishing the HBs between N439 or S443 and D104_H_ as well as the electrostatic interactions between K444 and D104_H_ (**Table S4** and **SI excel sheet**). Importantly, the loss/reduce of REGN10987 binding to Omicron subvariants is attributed largely to altering the interaction profile of RBD’s loop 1 (L1) (i.e., residues 441 to Y451) due to N440K and Q498R as well as G446S of BA.1. Specifically, Q498R loss the interactions with W99_L_ and D104_H_ and inducing internal steric clashes with S443, K444, V445, G/S446, P499, from RBD. These clashes have a variety of consequences, including interrupting the hydrophobic contacts between RBD residues (i.e., K444 to Y449 and P499) and W99_L_, A33_H_, Y35_H_, W47_H_, V50_H_, Y53_H_, Y59_H_, Y105_H_ from REGN10987. When G446 is mutated to S446 in BA.1, the HB with W99_L_ is missed and the interaction with W47_H_ is lost (**Figure 7**). Besides, the interactions of the conserved RBD residues are impacted especially R346 (**Table S4** and **SI excel sheet**).

#### S309

Our finding revealed that only G339D, S371F, and N440K Omicron mutations can directly pair with S309. G339D forms HBs with Y32_H_ and Y100_H_ and strengthens the pair with L110_H_. However, the self-interaction energy of D339 between its atoms is unfavorably large compared to G399 due to its large sidechain. Thus, the gain energy from pairing D339 with S309 is inadequate to offset the large intra-interaction, leading to a reduction in the overall energy from this substitution. N440K exhibits this behavior as well. S371L/F forms one very weak pair with Y50_L_. The surrounding conserved residues have a mixed response in terms of binding enhancement or reduction (**Table S4**). The Glycan forms substantial pairs with S309 and behaves differently in each case, which is too complicated to mention here. Again, we only recall their net difference per-residue trends that have a negative impact on S309 binding of BA.2 and BA.4/5 compared to RBD WT.

#### S2X259

S371L/F mutation reduces S2X259 binding by abolishing the HB with S55_H_ and ignoring the pair with W107_H_ (**Figure 7**). It may also have an impact on the interaction of adjacent unmutated residues, such as N370 with M54_H_, S55_H_, and, most importantly, K74_H_. The opposite happens with S373F, in which changing the hydroxymethyl group of serine to the imino ring of proline involves more hydrophobically interactions with CDRH2, primarily with I52_H_ and M57_H_, suggesting a reinforcement in the binding. However, the interaction of the conserved residue A372 with CDRH2, particularly S55_H_ to M57_H_, is adversely affected by either S371L/F or S373P mutations (**Table S4**). Likewise, S375F keeps the same HB with W107_H_ in addition to providing more hydrophobic interactions with CDRL3 notably with Y93_L_, P100_L_ and W102_L_. However, S375F could sterically clash within the internal RBD residues as well as with the S2X259 residues like M57_H_ and N59_H_. N501Y and Y505H have no impact due to maintaining weak contact with CDRL3.

In BA.2 and BA.4/5, R408S can primarily account for the significant reduction in the binding S2X259 due to revoking the attractive electrostatic and/or HB interactions with G32_L_, D34_L_, and D110_H_ (**Figure 7**). However, a salt-bridge can be formed between the preserved residue K378 with D109_H_ to counterbalance such charge interaction losing (**Table S4** and **SI excel sheet**). Unlike the positive charge R408 side chain in WT or BA.1, the small S408 side chain and its neutral charge do not interface or repel K378 in BA.2 and BA.4/5, facilitating this salt-bridge formation. T376A weakens the S2X259 binding even further due to the lack of either O…OH HB or the hydrophobic contacts with W107_H_, as well as reducing the interaction with G108_H_. Furthermore, D405N disrupts the HB networks with Y33_L_ and S95_L_, implying a reduction in S2X259 binding. At first glance, this appears to contradict what was demonstrated by the enhancement binding of this mutation using per-residue BFE decomposition (**Figure 6**). In fact, the total effect should be estimated rather than the local effect, so the net difference in per-residue BFE for all residues involved in interactions with this mutation should be considered. In other words, the net difference per-residue BFE of D405N, Y33_L_, and S95_L_ should be added together, which is consistent with our observation that D405N could hinder its binding.

Again, Omicron sublineages markedly reduce the strength and quantity of HBs of the mAb-RBD complex compared to their WTs (see **SI excel sheet**). Both the HB of the local mutation site and the nearest-local RBD sites can be disrupted.

## 3. Conclusion

On the basis of all-atom explicitly solvated MD simulations and comprehensive BFE analyses, we present a systematic investigation into antigenic characteristics of RBD Omicron subvariants, BA.1, BA.2 and BA.4/5, on the ten mAbs that target nearly all known RBD epitopes. The current study provides clear evidence that the binding of the majority of mAbs is adversely influenced by Omicron subvariants with exception of S309 and S2X259 against BA.1. The electrostatic interaction has been assigned as the primary physical factor that assists Omicron in evading the majority of mAb. The key Omicron mutations that confer antibody resistance has been also identified for each mAb. Furthermore, the structural and thermodynamic basis for these escape Omicron mutations has been thoroughly determined, and our findings show that the main source of the antibody evasion is the loss of ionic and hydrogen interactions between Omicron RBDs and mAbs. This research sheds light on the molecular mechanism by which mutations in the RBD Omicron subvariants affect the mAb-RBD recognition process. Additionally, this study paves the route for the development of effective antibody therapy against current or future SARS-CoV-2 variants.

## Supporting information

SI excel sheet

## ASSOCIATED CONTENT

### Supporting Information

Additional methods details and a full description of the procedure developed to create models. Table S1 of the atomic detailed of the solvated mAb-RBD models. Table S2 of the comparison of the calculated BFE in this study with experimental data. Table S3 of comparison of the RBD epitope identified in this study to those identified structurally. Table S4 of the AA-AA pair interactions that impact by Omicron mutations. Figures for all RMSDs vs time step and RMSFs of mAb-RBD complex, the dynamic conformations projected of two principal components of only RBD or entire mAb-RBD complex, scatter correlation plots of BFE, and RBD Omicron mutations impact on the interaction spectrum based on the per-residue analysis.

Additional excel sheet file for all HBs between RBD and mAb in all cases and models.

## AUTHOR INFORMATION

### Author Contributions

BJ and WC conceived the project. BJ performed the calculations and made most of the figures. BJ and WC drafted the paper with inputs from PA and RP. All authors participated in the discussion and interpretation of the results. All authors edited and proofread the final manuscript.

## ACKNOWLEDGMENT

This research used the resources of the National Energy Research Scientific Computing Center (NERSC), a DOE Office of Science User Facility supported by the Office of Science of the U.S. Department of Energy by U.S. Department of Energy under Contract No. DE-AC03-76SF00098, DE-AC02-05CH11231 using NERSC award NERSC DDR-ERCAP0023727, and the Research Computing Support Services (RCSS) of the University of Missouri System. We would like to thank Richard Gerber.

