## Supplementary material for "Impact of BA.1, BA.2, and BA.4/BA.5 Omicron Mutations on Therapeutic Monoclonal Antibodies": Table S1, Table S2, Table S3, Table S4, Figure S1, Figure S2, Figure S3, Figure S4, Figure S5, Figure S6, Figure S7

**1. Computational Modeling and Methods**

**1.1 Preparation of mAb-RBD structures with Omicron subvariant mutations**

Since several of the mAb-RBD complex structures of Omicron subvariants have yet to be resolved, we developed a procedure for creating initial structural models having all RBD Omicron mutations as described below.

i) *Building mAb structure.* Apart from providing valuable mAb-RBD structures using cryo-EM or X-ray experimental techniques, they are unable to resolve the highly flexible mAb regions (loops) due to their resolution limits.^1^ So, to create complete mAb, all non-terminal missing antibody residues for heavy and light chains (HC and LC) were modeled using the Modeller tool,^2^ which was implemented via the Chimera graphical interface.^3^ Here, this resulting antibody structure will remain unchanged when making their four mAb-RBD complexes. In other words, the RBDs of these complexes differ because they have substitutions at specific sites that characterize Omicron versions from WT. Note the PDBs for mAb will be explained below in point (iii).

ii) *Producing glycosylated RBD structure.* This step was to generate four glycosylated RBD structures (WT, BA.1, BA.2, or BA.4/5) with the same number of residues and glycan linked to its N343 position in all cases. In this regard, the individual WT RBD from ID: 7C01-Chain A was extracted and prepared for using in all WT complexes.^4^ It contains 195 residues from T333 to P527. Similarly, we selected Omicron BA.1 RBD alone from ID:7WBP of RBD-ACE2 complex,^5^ which has the same 195 AAs as well. This BA.1 RBD was used to produce BA.2 and BA.4/5 RBDs. As mentioned early, BA.2 RBD shares 12 mutations with BA.1 RBD and thus kept unchanged. While the other four (L371, A376, N405, and S408) mutations of BA.2 were formed using the Dunbrack backbone-dependent rotamer library,^6^ and are based on the same conformations of BA.2 RBD from ID:7ZF7 that was recently released.^7^ The reasons for not directly using BA.2 RBD of 7ZF7 are that it has lower resolution than BA.1 RBD of 7WBP (3.46 vs 3 Å) and one AA missing at its terminal. Next, the obtained BA.2 RBD was employed to generate BA.4/5 RBD using the same protocol. Specifically, the 15 BA.2-like mutations were unchanged, while L452 and F486 or R493 were substituted by R452, V486, and Q493 respectively. Because BA.4/5 does not have a mutation at 493, we returned R493 of BA.2 to its original WT or Q493. As a result, we made four RBDs with the same residue numbers that mimic the real experimental structures.

Owing to the important role of N-linked glycan in facilitating immune evasion by shielding certain epitopes from antibody binding,^8^ it should be included in our MD modeling. Therefore, the N-glycan was appropriately connected to N343 of the RBD using the protocol developed by Thomson et al.^9^ In brief, the FA2G2 glycosylation pattern was used to define and represent the structure of the N-glycan at N343 of RBD according to Watanabe et al.,^8^ Shajahan et al.,^10^ and Harbison et al. studies.^11^ Thomson and coworkers make the PDB FA2G2 structure file available online at <https://github.com/choderalab/rbd-ab-contact-analysis/tree/main/01_system_preparation/representative_glycan_structures>, which we used. Then, the N-Acetylglucosamine (NAG) of FA2G2 was aligned with NAG of our constructed RBDs (WT, BA.1, ect.) using a structure comparison tool in the UCSF Chimera software.^12^ Next, we deleted the NAG of RBD and replaced them by whole FA2G2. Further steps like relax clashes and covalent connectivity within the FA2G2 structure will describe in point (iv). The glycosylated RBDs are now ready to reassemble with the mAb Fab system, as shown in the next step.

iii) *Reassembling mAb with the relevant glycosylated RBD to form the mAb-RBD complex.* First, the available complex structures for mAb with WT RBD were used as a template to produce the related complexes with Omicron subvariant mutations. The PDB IDs for these template structures are 7R6X (S309_WT_),^13^ 7L7D (COV2-2196_WT_),^14^ 7L7E (COV2-2130_WT_),^14^ 6XDG (REGN10933_WT_ or REGN10987_WT_),^15^ 7KMG (LY-CoV555_WT_),^16^ 7C01 (LY-CoV016_WT_),^4^ 7CM4 (CT-P59_WT_),^17^ 7BWJ (BRII-196_WT_),^18^ and 7M7W (S2X250_WT_)^13^ (See **Table S1** for more details). Here for simplicity and distinction, we proposed the mAb_X_ symbol for each unique model, where mAb stands for the mAb Fab-RBD complex and subscript represents the different RBD cases (X: WT, BA.1, BA.2 or BA.4/5). For instance, we adopted this symbol to write S309_WT_ instead of S309 Fab-RBD_WT_.

Now, the glycosylated RBD from point (ii) was combined with the mAb from point (i) by alignment with its corresponding WT complex using UCSF Chimera, and then the original compartments of the complex were deleted. This process was repeated four times, once for each glycosylated RBD case, to yield four glycosylated complexes for each unique mAb.

iv) *Parametrization and Solvation of glycosylated mAb-RBD system.* The AmberTools18 tleap program was employed to prepare all solvated complexes for MD simulation,^19^ beginning with reading the mAb_X_ and adding hydrogen atoms, and then connecting all covalent bonds within the FA2G2 glycan structure and all RBD and mAb disulfide bonds. Here, the Amber ff14SB^20^ and GLYCAM 06j-1^21^ force fields were used to parameterize the inter- and intramolecular interactions of proteins (RBD and mAb) and glycan respectively. Subsequently, the glycosylated mAb-RBD complex was explicitly solvated with TIP3P water model in a periodic water box of 30000 water molecules.^22^ To neutralize the solvated system and mimic the physiological salt concentration of 0.15 M, the appropriate number of Na^+^ and Cl^-^ ions were randomly added using the Li/Merz ion parameters of monovalent ions for the TIP3P water model.^23^

All 40 solvated models of the glycosylated mAb-RBD complex in WT or Omicron variants are outlined in **Table S1** of SI.

**1.2 Molecular Dynamics (MD) Simulation and Binding Free Energy (BFE) Calculation**

All-atom MD simulations for each mAb_X_ were performed in an explicit solvent with periodic boundaries using the AMBER 18 software.^24^ This was followed by end-point binding free energy (BFE) calculations with molecular mechanics Generalized Born surface area (MM-GBSA) method to provide a complete binding profile.^25–28^ We implemented the same MD and BFE protocols that had previously been successfully applied to other systems,^29–31^ but with certain modifications. In briefly, the MD steps began with six stages of energy minimization with 5000-cycle steepest descent and 5000-cycle conjugate gradient to remove bad crashes. In the first five stages, a range of restraint force constants of 500, 250, 100, 10, and 1 kcal/mol-Å^2^ on the solute were applied sequentially, while no constraint was imposed in the sixth stage. Then, the system was incrementally heated from 0 to 310 K for 310 picoseconds (ps) using the NVT ensemble with a weak restraint of 10 kcal/mol-Å^2^ on the solute. Finally, two independent NTP MD simulations along 200 ns (the cumulative simulation time was 16 μs, i.e., 2 × 200 ns × 40 MD models) were performed after the system was equilibrated for 5 ns without restraint under NPT ensemble (310 K and 1 bar pressure). The following parameter settings were employed during the MD simulations: the particle mesh Ewald (PME) scheme for long-range electrostatic interactions;^32^ Langevin dynamics for temperature scaling; 2 ps as the pressure relaxation time; and 10 Å cutoff for the Lennard-Jones and direct space PME. All bonds containing hydrogen atoms were constrained using the SHAKE algorithm,^33^ allowing an integration time step of 2 fs to be employed. PMEMD.CUDA module of AMBER 18 was used as the simulation engine.^34^

Subsequently, the AMBER MM-PBSA.py module^35^ was implemented to compute BFE and provide the thermodynamic profile using the MM-GBSA methodology based on 500 snapshots extracted over the course of 200 ns MD simulations and following input parameters: igb=2 for GB model; 0.0072 kcal/mol-Å2 and zero for surface tension coefficient and offset correction, respectively, to calculate nonpolar solvation free energy; 78.15 and 1 for exterior and interior dielectric constants respectively; 0.15 M for salt concentration; and quasi-harmonic approximation to estimate entropic contribution based on only 5 snapshots along whole simulation. To identify the role and the nature of interaction for each individual Omicron mutation, the *Per-residue* and *pairwise BFE decompositions* were conducted.

**Table S1.** Atomic detailed of the complex between the monoclonal antibody (mAb) and glycosylated receptor binding domain (RBD) of spike protein for the wild-type strain and three Omicron subvariants (BA.1, BA.2 and BA.4/5). H and L represent the heavy and light chains of the fragment antigen binding (Fab) region of mAb. mAb_X_ is a proposed symbol to distinguish different models, where mAb stands for the mAb Fab-RBD complex and the subscript denotes the various RBD cases (X: WT, BA.1, BA.2, or BA.4/5).

| **Details of mAb** | | **Glycosylated RBD (atom number)** | **Name of the mAb-RBD model** | **Water molecules** | **Ions (Na^+^+Cl^-^)** | **Total number of atoms** |
| --- | --- | --- | --- | --- | --- | --- |
| **Amubarvimab (BRII-196)** | **Class :1**  PDB:7BWJ  H residue:1-229  L residue:1-213  Atom number: 6483  IGHV Gene: IGHV4-38-2  Brii Biosciences* | WT (3240) | BRII-196_WT_ | 30000 | 107+118 | 99948 |
|  |  | BA.1 (3296) | BRII-196_BA.1_ | 30000 | 107+121 | 100007 |
|  |  | BA.2 (3274) | BRII-196_BA.2_ | 30000 | 107+121 | 99985 |
|  |  | BA.4/5 (3268) | BRII-196_BA.4/5_ | 30000 | 107+121 | 99979 |
| **Regdanvimab (CT-P59)** | **Class:1**  PDB: 7CM4  H residue:1-231  L residue:1-214  Atom number: 6622  IGHV Gene: IGHV2-70  Celltrion* | WT (3240) | CT-P59_WT_ | 30000 | 107+116 | 100085 |
|  |  | BA.1 (3296) | CT-P59_BA.1_ | 30000 | 107+119 | 100144 |
|  |  | BA.2 (3274) | CT-P59_BA.2_ | 30000 | 107+119 | 100122 |
|  |  | BA.4/5 (3268) | CT-P59_BA.4/5_ | 30000 | 107+119 | 100116 |
| **Etesevimab (LY-CoV016)** | **Class :1**  PDB: 7C01  H residue: 1-218  L residue: 1-215  Atom number: 6477  IGHV Gene: IGHV3-66  Eli Lilly* | WT (3240) | LY-CoV016_WT_ | 30000 | 107+112 | 99936 |
|  |  | BA.1 (3296) | LY-CoV016_BA.1_ | 30000 | 107+115 | 99995 |
|  |  | BA.2 (3274) | LY-CoV016_BA.2_ | 30000 | 107+115 | 99973 |
|  |  | BA.4/5 (3268) | LY-CoV016_BA.4/5_ | 30000 | 107+115 | 99967 |
| **Bamlanivimab (LY-CoV555)** | **Class :2**  PDB: 7KMG  H residue: 1-228  L residue: 1-212  Atom number: 6624  IGHV Gene: IGHV3-69  Eli Lilly* | WT (3240) | LY-CoV555_WT_ | 30000 | 107+118 | 100089 |
|  |  | BA.1 (3296) | LY-CoV555_BA.1_ | 30000 | 107+121 | 100148 |
|  |  | BA.2 (3274) | LY-CoV555_BA.2_ | 30000 | 107+121 | 100126 |
|  |  | BA.4/5 (3268) | LY-CoV555_BA.4/5_ | 30000 | 107+121 | 100120 |
| **Tixagevimab (COV2-2196)** | **Class :2**  PDB: 7L7D  H residue: 1-225  L residue: 1-215  Atom number: 6584  IGHV Gene: IGHV1-58  AstraZeneca* | WT (3240) | COV2-2196_WT_ | 30000 | 107+116 | 100047 |
|  |  | BA.1 (3296) | COV2-2196_BA.1_ | 30000 | 107+119 | 100106 |
|  |  | BA.2 (3274) | COV2-2196_BA.2_ | 30000 | 107+119 | 100084 |
|  |  | BA.4/5 (3268) | COV2-2196_BA.4/5_ | 30000 | 107+119 | 100078 |
| **Casirivimab (REGN10933)** | **Class :2**  PDB: 6XDG  H residue: 1-220  L residue: 1-214  Atom number: 6500  IGHV Gene: IGHV3-11  Regeneron* | WT (3240) | REGN10933_WT_ | 30000 | 107+111 | 99958 |
|  |  | BA.1 (3296) | REGN10933_BA.1_ | 30000 | 107+114 | 100017 |
|  |  | BA.2 (3274) | REGN10933_BA.2_ | 30000 | 107+114 | 99995 |
|  |  | BA.4/5 (3268) | REGN10933_BA.4/5_ | 30000 | 107+114 | 99989 |
| **Imdevimab (REGN10987)** | **Class :3**  PDB: 6XDG  H residue: 1-220  L residue: 3-213  Atom number: 6351  IGHV Gene: IGHV3-30  Regeneron* | WT (3240) | REGN10987_WT_ | 30000 | 107+115 | 99813 |
|  |  | BA.1 (3296) | REGN10987_BA.1_ | 30000 | 107+118 | 99872 |
|  |  | BA.2 (3274) | REGN10987_BA.2_ | 30000 | 107+118 | 99850 |
|  |  | BA.4/5 (3268) | REGN10987_BA.4/5_ | 30000 | 107+118 | 99844 |
| **Cilgavimab (COV2-2130)** | **Class :3**  PDB: 7L7E  H residue: 1-236  L residue: 1-219  Atom number: 6838  IGHV Gene: IGHV3-15  AstraZeneca* | WT (3240) | COV2-2130_WT_ | 30000 | 107+114 | 100299 |
|  |  | BA.1 (3296) | COV2-2130_BA.1_ | 30000 | 107+117 | 100358 |
|  |  | BA.2 (3274) | COV2-2130_BA.2_ | 30000 | 107+117 | 100336 |
|  |  | BA.4/5 (3268) | COV2-2130_BA.4/5_ | 30000 | 107+117 | 100330 |
| **Sotrovimab (S309)** | **Class :3**  PDB: 7R6X  H residue: 1-228  L residue: 1-213  Atom number: 6596  IGHV Gene: IGHV1-30  Vir Biotechnology/ GlaxoSmithKline* | WT (3240) | S309_WT_ | 30000 | 107+115 | 100058 |
|  |  | BA.1 (3296) | S309_BA.1_ | 30000 | 107+118 | 100117 |
|  |  | BA.2 (3274) | S309_BA.2_ | 30000 | 107+118 | 100095 |
|  |  | BA.4/5 (3268) | S309_BA.4/5_ | 30000 | 107+118 | 100089 |
| **(S2X259)** | **Class :4**  PDB: 7M7W  H residue: 2-228  L residue: 2-217  Atom number: 6494  IGHV Gene: IGHV1-69 | WT (3240) | SX259_WT_ | 30000 | 107+113 | 99954 |
|  |  | BA.1 (3296) | SX259_BA.1_ | 30000 | 107+116 | 100013 |
|  |  | BA.2 (3274) | SX259_BA.2_ | 30000 | 107+116 | 99991 |
|  |  | BA.4/5 (3268) | SX259_BA.4/5_ | 30000 | 107+116 | 99985 |

* is the developer company

**Table S2.** Comparison of the predicted binding free energy $(\Delta G_{Cal})$ using MM-GBSA method with the experimental binding data for the wild-type (WT) mAb-RBD complexes ^4,13–18,36^. The experimental measured values of the equilibrium dissociation constant (KD) are converted to $\Delta G_{exp}$ at T = 310K using $\Delta G_{Exper}=RTln(K_{D})$.

| **WT Model** | **Dissociation Constant *K_D_* (nM)^#^** | $\boldsymbol{\Delta}\boldsymbol{G}_{\boldsymbol{Exper}}$ **(kcal/mol)** | $\boldsymbol{\Delta}\boldsymbol{G}_{\boldsymbol{Cal}}$ **(kcal/mol)** |
| --- | --- | --- | --- |
| **BRII-196** | 5.14 | -11.78 | -4.83 ± 0.23 |
| **CT-P59** | 0.027 or 0.251 | -15.01 or -13.64 | -14.01 ± 0.3 |
| **LY-CoV016** | 2.49 | -12.22 | -21.15 ± 0.4 |
| **LY-CoV555** | 0.024 | -15.09 | -16.14 ± 0.3 |
| **COV2-2196** | 0.253 | -13.63 | -13.03 ± 0.5 |
| **COV2-2130** | <0.01 | <-17.05 | -17.84 ± 0.31 |
| **REGN10933** | 1.15 | -12.7 | -12.94 ± 0.42 |
| **REGN10987** | 45.2 | -10.43 | -11.47 ± 0.4 |
| **S309** | <0.01 or 0.2 | <-17.05 or -13.8 | -37.54 ± 0.6 |
| **S2X259** | 0.08 | -14.35 | -23.33 ± 0.5 |

**Table S3.** Comparison of the RBD epitope for each mAb using structural basis from IEDB database and thermodynamic and energetic basis using MM-GBSA method. Blue and Red: Predicated site based on MD study. Blue represents those who contributed to Per-Residue BFE and made fewer than two pairs with mAb, while red represents those who made more than two pairs with mAb. Bold: The sites where Omicron subvariant mutations occur.

| mAb (PDB) | Method | RBD Epitope |
| --- | --- | --- |
| BRII-196 (7BWJ) | Structure | R346, K444, **G446**, G447, N448, Y449, N450, **L452**, V483, **E484**, G485, F490, S494 |
|  | Our MD | R346, K444, **G446**, G447, N448, Y449, N450, **L452**, V483, **E484**, G485, F490, S494 |
|  |  | V445, T470, I472, **F486**, L492, **Q493** |
| CT-P59 (7CM4) | Structure | R403, **K417**, Y449, N450, **L452**, Y453, L455, F456, **E484**, G485, **F486**, Y489, F490, L492, **Q493**, S494, Y495, **Y505** |
|  | Our MD | R403, **K417**, Y449, N450, **L452**, Y453, L455, F456, **E484**, G485, **F486**, Y489, F490, L492, **Q493**, S494, Y495, **Y505** |
|  |  | Y451, V483, C488, P491 |
| LY-CoV016  (7C01) | Structure | R403, **D405**, E406, **R408**, Q409, T415, G416, **K417**, D420, Y421, L455, F456, R457, K458, N460, Y473, Q474, A475, G476, **S477**, **F486**, N487, Y489, **Q493**, Y495, G502, **Y505** |
|  | Our MD | R403, **D405**, E406, **R408**, Q409, T415, G416, **K417**, D420, Y421, L455, F456, R457, K458, N460, Y473, Q474, A475, G476, **S477**, **F486**, N487, Y489, **Q493**, Y495, G502, **Y505** |
|  |  | K424, S459, **T478** |
| LY-CoV555  (7C01) | Structure | Y351, Y449, L455, T470, N481, G482, V483, **E484**, G485, **F486**, C488, Y489, F490, L492, **Q493**, S494 |
|  | Our MD | Y351, Y449, L455, T470, N481, G482, V483, **E484**, G485, **F486**, C488, Y489, F490, L492, **Q493**, S494 |
|  |  | N450, **L452**, F456, I472 |
| COV2-2196 (7L7D or 7L7E) | Structure | L455, A475, G476, **S477**, **T478**, **E484**, G485, **F486**, N487, C488, Y489, **Q493** |
|  | Our MD | L455, A475, G476, **S477**, **T478**, **E484**, G485, **F486**, N487, C488, Y489, **Q493** |
|  |  | **K417**, Y449, F456, Y473, P479, F490, **Y505** |
| COV2-2130 (7L7E) | Structure | R346, N439, **N440**, L441, S443, K444, V445, **G446**, Y449, N450, **L452**, **E484**, F490, **Q493**, S494, P499 |
|  | Our MD | R346, N439, **N440**, L441, S443, K444, V445, **G446**, Y449, N450, **L452**, **E484**, F490, **Q493**, S494, P499 |
|  |  | T345, G447, N448, L492 |
| S309  (7R6X) | Structure | N334, L335, P337, **G339**, E340, N343, A344, T345, R346, K356, R357, S359, C361, L441 |
|  | Our MD | N334, L335, P337, **G339**, E340, N343, A344, T345, R346, K356, R357, S359, C361, L441 |
|  |  | C336, V341, I358, **N440**, K444, V445, R509 |
| RENG10933 (6XDG) | Structure | **K417**, Y453, L455, F456, **E484**, G485, **F486**, N487, C488, Y489, **Q493** |
|  | Our MD | **K417**, Y453, L455, F456, **E484**, G485, **F486**, N487, C488, Y489, **Q493** |
|  |  | R403, E406, Y421, Y473, A475, G476, **S477**, **T478**, **Q498**, **Y505** |
| RENG10987 (6XDG) | Structure | R346, **N440**, L441, K444, V445, **G446**, N448, Y449, **Q498** |
|  | Our MD | R346, **N440**, L441, K444, V445, **G446**, N448, Y449, **Q498** |
|  |  | T345, N439, S443, N450, Y451, F497, P499 , T500 |
| S2X259 (7M7W) | Structure | Y369, N370, **S371**, A372, F374, **S375**, **T376**, F377, K378, C379, Y380, V382, S383, P384, T385, G404, **D405**, **R408**, T500, **N501**, G502, V503, G504, Q506 |
|  | Our MD | Y369, N370, **S371**, A372, F374, **S375**, **T376**, F377, K378, C379, Y380, V382, S383, P384, T385, G404, **D405**, **R408**, T500, **N501**, G502, V503, G504, Q506 |
|  |  | **S373**, G381, V407 |

**Table S4.** All possible RBD:mAb pairs that impact due to the RBD Omicron mutations using pairwise BFE decomposition of MM-GBSA method. First, all potential pairs in all Omicron and WT models are accounted and then the change in the pairwise BFE decomposition for each pair with respect to its WT pair is calculated using ${\Delta\Delta G}_{\mathrm{Pairwise}}={(\Delta\Delta G}_{\mathrm{Pairwise}})_{BA.i}-{(\Delta\Delta G}_{\mathrm{Pairwise}})_{WT}$ where *BA.i* can be BA.1, BA.2 or BA.4/5. Energy differences larger than -0.15 or less than 0.15 kcal/mol are ignored. The RBD:mAb pairs at the mutated site is highlighted in yellow. The mAb light chair residues are labeled with asterisk to distinguish from heavy chain residues. ΔBA.1, ΔBA.2, or ΔBA.4 represent the change in the pairwise BFE with respect to WT. All units in kcal/mol.

| **Table S4: BRII-196** | | | |  |  |  |  |  |
| --- | --- | --- | --- | --- | --- | --- | --- | --- |
| RBD:mAb pair | ΔBA.1 | ΔBA.2 | ΔBA.4 |  |  |  |  |  |
| K444:S30 | -0.99 | 0.31 | -0.84 |  |  |  |  |  |
| V445:G26 | 0.40 | 0.20 | 0.09 |  |  |  |  |  |
| V445:Y27 | 0.72 | 0.41 | -0.14 |  |  |  |  |  |
| V445:S28 | -0.25 | -0.11 | -1.11 |  |  |  |  |  |
| V445:S31 | -0.08 | -0.02 | -0.67 |  |  |  |  |  |
| G446S:Y27 | -1.13 |  |  |  |  |  |  |  |
| G446S:S28 | -1.79 |  |  |  |  |  |  |  |
| G446S:S31 | -0.89 |  |  |  |  |  |  |  |
| G447:Y27 | 1.93 | 0.78 | 1.03 |  |  |  |  |  |
| G447:S31 | -0.24 | -0.12 | -0.30 |  |  |  |  |  |
| Y449:S31 | 0.12 | 0.03 | 0.52 |  |  |  |  |  |
| Y449:G32 | 0.01 | 0.06 | 0.18 |  |  |  |  |  |
| Y449:Y33 | 0.32 | 0.13 | 0.04 |  |  |  |  |  |
| Y449:G102 | -0.19 | 0.03 | 0.10 |  |  |  |  |  |
| Y449:I103 | -0.52 | -0.36 | -0.24 |  |  |  |  |  |
| N450:S30 | 0.90 | 0.31 | 0.85 |  |  |  |  |  |
| N450:S31 | 0.42 | 0.16 | 0.20 |  |  |  |  |  |
| N450:G32 | 0.22 | 0.04 | 0.10 |  |  |  |  |  |
| N450:Y53 | -0.25 | -0.44 | 0.12 |  |  |  |  |  |
| N450:H54 | 0.58 | -0.11 | 0.58 |  |  |  |  |  |
| L452R:I103 |  |  | 0.15 |  |  |  |  |  |
| L452R:V105 |  |  | -0.23 |  |  |  |  |  |
| I472:P107 | 0.37 | 0.20 | 0.31 |  |  |  |  |  |
| N481:Y32* | -0.16 | -0.43 | 0.02 |  |  |  |  |  |
| N481:N33* | -1.79 | -1.33 | -0.12 |  |  |  |  |  |
| G482:Y34* | -0.59 | -0.83 | -0.15 |  |  |  |  |  |
| G482:R112 | -0.23 | -0.36 | -0.03 |  |  |  |  |  |
| V483:G31* | 0.38 | 0.27 | 0.23 |  |  |  |  |  |
| V483:Y32* | 1.03 | 0.66 | 0.47 |  |  |  |  |  |
| V483:N33* | -0.77 | -0.64 | -0.26 |  |  |  |  |  |
| V483:E52* | -1.71 | -1.11 | -0.23 |  |  |  |  |  |
| E484A:V106 | 0.61 | 0.61 | 0.62 |  |  |  |  |  |
| E484A:P107 | 0.70 | 0.58 | 0.47 |  |  |  |  |  |
| E484A:R112 | 12.48 | 12.44 | 12.32 |  |  |  |  |  |
| E484A:Y32* | 0.38 | 0.33 | 0.31 |  |  |  |  |  |
| E484A:N33* | 2.32 | 1.86 | 1.55 |  |  |  |  |  |
| E484A:Y34* | 5.02 | 4.71 | 4.23 |  |  |  |  |  |
| E484A:E52* | -1.11 | -0.41 | -0.37 |  |  |  |  |  |
| E484A:K55* | -1.03 | -1.12 | -0.32 |  |  |  |  |  |
| G485:N33* | 0.87 | 0.70 | 0.53 |  |  |  |  |  |
| G485:K55* | -0.64 | -0.28 | -0.13 |  |  |  |  |  |
| F486V:N33* |  |  | 0.15 |  |  |  |  |  |
| F490:I103 | -0.12 | 0.04 | 0.30 |  |  |  |  |  |
| F490:V105 | 0.31 | 0.27 | 0.49 |  |  |  |  |  |
| F490:V106 | 0.67 | 0.56 | 1.06 |  |  |  |  |  |
| F490:P107 | 0.80 | 0.55 | 0.42 |  |  |  |  |  |
| Q493R:I103 | 0.32 | 0.37 |  |  |  |  |  |  |
| S494:I103 | 0.23 | 0.22 | 0.35 |  |  |  |  |  |
| **Table S4: CT-P59** | | | | | | | | |
| RBD:mAb pair | ΔBA.1 | ΔBA.2 | ΔBA.4 |  | RBD:mAb pair | ΔBA.1 | ΔBA.2 | ΔBA.4 |
| R403:D54 | 0.14 | 0.27 | 0.31 |  | F486V:P101 |  |  | -0.18 |
| R403:D56 | -2.42 | 6.97 | 9.08 |  | F486V:G102 |  |  | 0.33 |
| R403:D57 | -0.05 | 0.24 | 0.27 |  | F486V:Y111 |  |  | 1.58 |
| R403:N58 | -0.32 | 0.10 | 0.12 |  | F486V:Y113 |  |  | 1.54 |
| K417N:S32 | 1.21 | 1.21 | 1.22 |  | Y489:T31 | -0.16 | -0.02 | -0.75 |
| K417N:W55 | 0.02 | 0.22 | 0.29 |  | Y489:S32 | -1.58 | -0.01 | -1.66 |
| K417N:D56 | 0.06 | 0.41 | 0.41 |  | Y489:G33 | -0.79 | 0.06 | -0.59 |
| G446S:R105 | -0.59 |  |  |  | Y489:P101 | -0.41 | 0.11 | -0.07 |
| Y449:N58 | 0.03 | -0.95 | -1.75 |  | Y489:G102 | 0.30 | 0.27 | 0.64 |
| Y449:Y60 | 0.37 | -0.11 | -0.22 |  | Y489:L104 | 1.27 | 0.39 | 1.67 |
| Y449:R105 | 3.58 | 0.14 | -0.81 |  | Y489:Y106 | 0.34 | -0.30 | 0.37 |
| Y449:Y106 | -1.08 | -0.39 | -2.01 |  | Y489:Y111 | 0.96 | 0.07 | 1.23 |
| Y449:R107 | -1.05 | -0.24 | -0.65 |  | F490:L104 | -0.72 | -0.18 | -1.14 |
| Y449:N108 | -1.55 | 0.12 | 0.08 |  | F490:Y106 | 1.19 | 0.30 | 3.07 |
| Y449:Y110 | -0.52 | 0.35 | 0.42 |  | F490:R109 | 0.89 | 0.28 | 0.89 |
| N450:Y106 | 0.00 | -0.02 | -1.86 |  | F490:Y111 | 0.11 | -0.07 | -0.50 |
| N450:R107 | 2.95 | -0.58 | -1.80 |  | L492:R105 | 0.23 | 0.27 | 0.31 |
| L452R:Y106 |  |  | -3.11 |  | L492:Y106 | 0.22 | 0.52 | 1.68 |
| L452R:R107 |  |  | 1.84 |  | Q493R:D54 | -0.48 | -3.03 |  |
| Y453:W55 | 0.41 | 0.31 | 0.27 |  | Q493R:W55 | 1.02 | 0.30 |  |
| Y453:D56 | -2.94 | -0.33 | -0.19 |  | Q493R:D56 | -0.26 | -1.13 |  |
| Y453:R105 | -0.46 | 0.01 | 0.02 |  | Q493R:F103 | -0.64 | -0.07 |  |
| L455:S32 | 0.89 | 0.84 | 0.59 |  | Q493R:L104 | 0.66 | 0.73 |  |
| L455:W55 | 0.02 | 0.89 | 0.29 |  | Q493R:R105 | 1.19 | 4.03 |  |
| L455:D56 | -1.01 | 0.01 | -0.03 |  | Q493R:Y106 | -1.23 | -0.31 |  |
| L455:L104 | 0.60 | 0.70 | 0.71 |  | S494:R105 | -0.12 | 2.51 | 2.24 |
| F456:S32 | 0.31 | 1.03 | 0.30 |  | S494:Y106 | -0.29 | 0.49 | 0.73 |
| F456:W55 | -0.83 | -0.01 | -0.08 |  | S494:R107 | 0.36 | 0.21 | 0.52 |
| F456:L104 | 0.56 | 0.64 | 0.71 |  | Y495:R105 | -1.10 | 0.01 | 0.02 |
| T470:Y106 | 0.46 | 0.31 | 0.62 |  | G496S:R105 | -2.05 |  |  |
| Y473:S32 | -0.31 | 0.02 | 0.00 |  | Q498R:S94* | -0.24 | -0.01 | 0.00 |
| A475:S32 | -0.40 | 0.01 | 0.00 |  | Q498R:S95* | -0.27 | 0.00 | 0.00 |
| N481:Y50* | 0.00 | -0.01 | -0.54 |  | Q498R:S97* | -0.92 | -0.07 | 0.01 |
| N481:K54* | 0.05 | -0.05 | -1.11 |  | Q498R:D56 | -0.03 | -2.60 | -0.13 |
| V483:Y33* | 0.27 | -0.08 | -0.33 |  | Q498R:D57 | -0.02 | -0.71 | -0.07 |
| V483:Y50* | -0.01 | -0.03 | -0.93 |  | Q498R:N58 | 0.00 | -1.68 | -1.40 |
| V483:D51* | 0.00 | -0.06 | -1.02 |  | Q498R:K59 | -0.03 | -0.28 | -0.24 |
| V483:K54* | -0.10 | -0.15 | -0.65 |  | N501Y:D56 | 0.00 | -1.16 | -0.10 |
| V483:Y111 | -0.02 | -0.02 | -0.78 |  | N501Y:N58 | -0.11 | -0.57 | -0.31 |
| V483:Y113 | 0.00 | -0.01 | -0.70 |  | N501Y:R105 | -1.15 | -0.01 | -0.01 |
| E484A:Y33* | 3.35 | 2.00 | 3.64 |  | Y505H:D56 | 1.77 | 1.18 | 1.85 |
| E484A:R109 | 9.61 | 9.55 | 9.61 |  | Y505H:D57 | 0.94 | 1.31 | 1.56 |
| E484A:Y111 | 0.19 | 0.38 | -1.22 |  | Y505H:N58 | -0.15 | 0.74 | 0.97 |
| E484A:Y106 | 0.33 | 0.30 | 0.34 |  | Y505H:K59 | -0.10 | 0.30 | 0.32 |
| G485:Y33* | 0.62 | -0.03 | 0.79 |  |  |  |  |  |
| G485:D51* | 2.60 | 0.94 | 3.58 |  |  |  |  |  |
| G485:P101 | -0.08 | -0.04 | -0.88 |  |  |  |  |  |
| G485:G102 | -0.10 | -0.01 | -0.48 |  |  |  |  |  |
| G485:Y111 | 0.67 | 0.19 | 1.03 |  |  |  |  |  |
| G485:Y113 | -0.29 | -0.10 | -0.88 |  |  |  |  |  |
| F486V:Y50* |  |  | 1.52 |  |  |  |  |  |
| F486V:D51* |  |  | 3.40 |  |  |  |  |  |
| F486V:K54* |  |  | 0.40 |  |  |  |  |  |
| F486V:F27 |  |  | -0.30 |  |  |  |  |  |
| F486V:T31 |  |  | -0.30 |  |  |  |  |  |
| F486V:V34 |  |  | -0.38 |  |  |  |  |  |
| F486V:R99 |  |  | -0.79 |  |  |  |  |  |

| **Table S4: LY-CoV061** | | | | | | | | |
| --- | --- | --- | --- | --- | --- | --- | --- | --- |
| RBD:mAb pair | ΔBA.1 | ΔBA.2 | ΔBA.4 |  | RBD:mAb pair | ΔBA.1 | ΔBA.2 | ΔBA.4 |
| R403:Y32* | 0.33 | 1.73 | 0.46 |  | N501Y:S30* | -0.11 | -0.37 | -0.24 |
| R403:Y92* | -1.25 | 0.36 | -2.40 |  | N501Y:R31* | -0.32 | -0.12 | -0.23 |
| R403:T94* | 0.05 | -1.43 | -0.47 |  | N501Y:Y32* | -0.06 | -0.64 | -0.04 |
| R403:Y102 | 0.01 | -3.36 | 0.12 |  | N501Y:Y92* | 0.00 | -0.74 | -0.04 |
| D405N:Y92* |  | 2.58 | 2.15 |  | N501Y:Y102 | -0.19 | -0.75 | -0.01 |
| D405N:S93* |  | -0.73 | -0.47 |  | G502:S28* | 0.18 | 0.21 | 0.05 |
| D405N:T94* |  | -0.99 | -1.31 |  | G502:Y92* | 0.00 | -0.29 | -0.07 |
| D405N:P95* |  | -0.22 | -0.18 |  | Y505H:S28* | 0.66 | 0.91 | 0.90 |
| E406:Y92* | -1.98 | 0.06 | 0.13 |  | Y505H:I29* | 0.68 | 0.61 | 0.80 |
| E406:S93* | -0.99 | -0.50 | -0.22 |  | Y505H:S30* | 0.33 | 1.42 | 0.69 |
| E406:T94* | -1.50 | -0.53 | -1.78 |  | Y505H:R31* | 0.29 | 0.71 | 0.15 |
| R408S:D1* |  | 0.80 | 0.80 |  | Y505H:Y92* | 0.97 | -1.66 | 0.28 |
| R408S:Y92* |  | 0.22 | 0.22 |  |  |  |  |  |
| R408S:T94* |  | 1.87 | 1.72 |  |  |  |  |  |
| R408S:P95* |  | 0.50 | 0.46 |  |  |  |  |  |
| Q409:T94* | 0.38 | 0.73 | 0.37 |  |  |  |  |  |
| T415:S56 | -0.47 | -0.15 | -0.39 |  |  |  |  |  |
| K417N:Y33 | 1.26 | 1.05 | 1.29 |  |  |  |  |  |
| K417N:Y52 | 1.45 | 1.56 | 1.67 |  |  |  |  |  |
| K417N:P100 | 1.45 | 1.50 | 1.47 |  |  |  |  |  |
| K417N:G103 | -0.50 | -0.50 | -0.50 |  |  |  |  |  |
| K417N:D104 | 10.29 | 10.35 | 10.32 |  |  |  |  |  |
| K417N:T94* | -0.06 | -0.43 | -0.38 |  |  |  |  |  |
| D420:S56 | 0.35 | 1.59 | -0.16 |  |  |  |  |  |
| Y421:Y33 | -0.40 | -0.05 | -0.28 |  |  |  |  |  |
| Y421:Y52 | -0.17 | 0.46 | 0.17 |  |  |  |  |  |
| Y421:S53 | -0.18 | -0.42 | -0.15 |  |  |  |  |  |
| Y421:S56 | 0.06 | 0.25 | 0.26 |  |  |  |  |  |
| Y453:Y32* | 0.27 | 0.32 | 0.25 |  |  |  |  |  |
| L455:Y33 | 0.61 | 0.10 | 0.78 |  |  |  |  |  |
| L455:P100 | 0.05 | 0.30 | 0.24 |  |  |  |  |  |
| F456:P100 | 0.02 | 0.74 | 0.23 |  |  |  |  |  |
| R457:S53 | -0.34 | 1.01 | -0.30 |  |  |  |  |  |
| N460:G54 | 0.42 | 0.46 | 0.06 |  |  |  |  |  |
| N460:S56 | 0.49 | 1.23 | 0.82 |  |  |  |  |  |
| A475:R97 | 0.22 | 0.19 | 0.14 |  |  |  |  |  |
| S477N:E1 | 0.12 | 0.26 | 0.45 |  |  |  |  |  |
| S477N:G26 | -0.30 | -0.63 | -0.52 |  |  |  |  |  |
| S477N:F27 | -0.20 | -0.33 | -0.14 |  |  |  |  |  |
| S477N:T28 | -0.23 | -0.24 | -0.15 |  |  |  |  |  |
| T478K:E1 | -1.62 | -1.62 | -0.56 |  |  |  |  |  |
| T478K:G26 | 0.26 | 0.26 | 0.26 |  |  |  |  |  |
| F486V:E1 |  |  | 0.18 |  |  |  |  |  |
| F486V:V2 |  |  | 0.73 |  |  |  |  |  |
| F486V:F27 |  |  | 0.27 |  |  |  |  |  |
| F486V:R97 |  |  | 0.44 |  |  |  |  |  |
| N487:G26 | 0.20 | 0.23 | 0.30 |  |  |  |  |  |
| Y489:M101 | 0.48 | 0.88 | 0.07 |  |  |  |  |  |
| Q493R:P100 | 0.04 | -1.94 | 0.04 |  |  |  |  |  |
| Q493R:M101 | 0.46 | 0.41 | 0.46 |  |  |  |  |  |
| Q493R:Y102 | 1.77 | 2.45 | 1.77 |  |  |  |  |  |
| Q493R:Y32* | 0.21 | -0.31 | 0.21 |  |  |  |  |  |
| Y495:Y102 | -0.48 | -0.71 | -0.16 |  |  |  |  |  |

| **Table S4: LY-CoV555** | | | | | | | | |
| --- | --- | --- | --- | --- | --- | --- | --- | --- |
| RBD:mAb pair | ΔBA.1 | ΔBA.2 | ΔBA.4 |  | RBD:mAb pair | ΔBA.1 | ΔBA.2 | ΔBA.4 |
| G446S:T28 | 0.42 |  |  |  | F490:R50 | 0.21 | 0.27 | 0.31 |
| G446S:R104 | -0.22 |  |  |  | F490:I52 | 0.21 | 0.35 | 0.79 |
| Y449:S30 | 0.60 | 0.96 | 1.41 |  | F490:I57 | 0.34 | 0.37 | 0.38 |
| Y449:T28 | 0.63 | 0.18 | -0.05 |  | F490:Y101 | 0.72 | -0.10 | -0.03 |
| Y449:N31 | 0.21 | 0.39 | -0.40 |  | F490:E102 | 0.34 | -0.35 | -0.20 |
| Y449:I54 | 0.48 | 0.45 | 0.79 |  | Q493R:E102 | 2.18 | 1.62 | 0.31 |
| Y449:Y101 | -0.65 | 0.00 | 0.00 |  | Q493R:A103 | -0.47 | -0.20 | -0.75 |
| Y449:E102 | -1.25 | -0.01 | -0.03 |  | Q493R:R104 | 1.38 | 0.48 | -1.25 |
| N450:I54 | 0.20 | 0.06 | 0.61 |  | S494:I54 | 0.24 | 0.15 | 0.24 |
| N450:K74 | 0.04 | 0.53 | 1.28 |  | S494:E102 | 1.25 | -0.79 | -4.00 |
| L452R:N31 |  |  | -0.60 |  | S494:R104 | 1.13 | 0.85 | -1.27 |
| L452R:I54 |  |  | 0.29 |  | G496S:R104 | -0.51 |  |  |
| L452R:L55 |  |  | 0.23 |  |  |  |  |  |
| L452R:Y101 |  |  | -0.30 |  |  |  |  |  |
| L452R:E102 |  |  | -0.90 |  |  |  |  |  |
| F456:A103 | 0.56 | 0.25 | 0.15 |  |  |  |  |  |
| T470:I57 | 0.01 | 0.20 | 0.80 |  |  |  |  |  |
| T470:N59 | -0.76 | -0.01 | 0.02 |  |  |  |  |  |
| T478K:D1* | 0.05 | -0.08 | -0.53 |  |  |  |  |  |
| T478K:Y92* | 0.02 | 0.31 | -0.78 |  |  |  |  |  |
| T478K:S93* | -0.22 | -0.02 | -0.72 |  |  |  |  |  |
| T478K:T94* | -0.09 | -0.06 | -0.44 |  |  |  |  |  |
| G482:R50 | -0.45 | 0.01 | 0.89 |  |  |  |  |  |
| G482:N59 | 0.10 | 0.53 | 0.36 |  |  |  |  |  |
| V483:W47 | 0.20 | 0.48 | 0.48 |  |  |  |  |  |
| V483:R50 | -0.46 | -0.20 | -0.26 |  |  |  |  |  |
| V483:N59 | 0.12 | 0.85 | 0.11 |  |  |  |  |  |
| V483:T94* | 0.10 | 0.84 | 0.98 |  |  |  |  |  |
| V483:R96* | -0.53 | -0.51 | -0.18 |  |  |  |  |  |
| E484A:R50 | 14.51 | 14.56 | 14.01 |  |  |  |  |  |
| E484A:Y101 | 1.49 | 1.17 | 0.85 |  |  |  |  |  |
| E484A:Y110 | 0.91 | 0.81 | 1.33 |  |  |  |  |  |
| E484A:S93* | 0.18 | 0.20 | 0.31 |  |  |  |  |  |
| E484A:T94* | 0.30 | 0.37 | 0.37 |  |  |  |  |  |
| E484A:R96* | 8.39 | 10.07 | 10.87 |  |  |  |  |  |
| G485:Y106 | -0.12 | -0.14 | -0.55 |  |  |  |  |  |
| G485:Y110 | 1.19 | 0.51 | 0.88 |  |  |  |  |  |
| G485:Y32* | -0.37 | -0.29 | 0.62 |  |  |  |  |  |
| G485:S91* | 0.16 | 0.14 | 0.50 |  |  |  |  |  |
| G485:Y92* | 0.09 | 0.45 | 1.32 |  |  |  |  |  |
| G485:S93* | 0.09 | 0.26 | 0.40 |  |  |  |  |  |
| F486V:Y106 |  |  | -1.73 |  |  |  |  |  |
| F486V:Y109 |  |  | -0.68 |  |  |  |  |  |
| F486V:Y32* |  |  | 0.82 |  |  |  |  |  |
| F486V:S91* |  |  | -0.49 |  |  |  |  |  |
| F486V:R96* |  |  | -0.61 |  |  |  |  |  |
| F486V:Y92* |  |  | 3.25 |  |  |  |  |  |
| N487:Y32* | 0.34 | 0.33 | 0.26 |  |  |  |  |  |
| N487:Y92* | -0.01 | 0.01 | -0.27 |  |  |  |  |  |
| N487:Y106 | 0.18 | 0.04 | -1.24 |  |  |  |  |  |
| C488:Y110 | 0.39 | 0.32 | 0.51 |  |  |  |  |  |
| Y489:A103 | 0.64 | 0.15 | -0.09 |  |  |  |  |  |
| Y489:Y106 | 0.19 | -0.17 | -1.24 |  |  |  |  |  |

| **Table S4: COV2-2196** | | | | | | | | |
| --- | --- | --- | --- | --- | --- | --- | --- | --- |
| RBD:mAb pair | ΔBA.1 | ΔBA.2 | ΔBA.4 |  | RBD:mAb pair | ΔBA.1 | ΔBA.2 | ΔBA.4 |
| K417N:M30 | 0.38 | 0.30 | 0.48 |  | F490:S55 | 0.35 | 0.55 | 0.61 |
| K417N:S103 | 1.19 | 1.18 | 1.19 |  | L492:S55 | 0.21 | 0.41 | 0.34 |
| K417N:I104 | 0.23 | 0.23 | 0.24 |  | Q493R:M30 | 0.59 | 0.83 |  |
| Y421:I104 | 0.22 | 0.20 | 0.19 |  | Q493R:G54 | 0.94 | 1.74 |  |
| Y449:R72 | 0.37 | 0.38 | 0.38 |  | Q493R:S55 | -0.64 | -0.55 |  |
| Y449:M74 | 0.21 | 0.21 | 0.21 |  | Q493R:G56 | -0.17 | -0.25 |  |
| Y453:M30 | 0.19 | 0.26 | 0.28 |  | Q493R:N57 | -0.34 | -1.39 |  |
| L455:M30 | 0.35 | 0.53 | 0.72 |  | Y505H:S75 | 0.54 | 0.65 | 0.65 |
| L455:S55 | -0.32 | -0.64 | -0.64 |  | Y505H:M74 | 0.15 | 0.24 | 0.24 |
| F456:S31 | 0.32 | 0.21 | 0.38 |  |  |  |  |  |
| F456:V52 | -0.06 | -0.25 | -0.24 |  |  |  |  |  |
| F456:G54 | -0.04 | -0.44 | -0.28 |  |  |  |  |  |
| F456:S55 | -0.07 | -0.36 | -0.27 |  |  |  |  |  |
| F456:I104 | 0.17 | 0.34 | 0.30 |  |  |  |  |  |
| K458:I104 | -0.94 | -1.49 | -0.20 |  |  |  |  |  |
| K458:S105 | -0.23 | -0.28 | -0.10 |  |  |  |  |  |
| Y473:I104 | 0.97 | 0.68 | 0.51 |  |  |  |  |  |
| Q474:D108 | 0.20 | 0.19 | 0.13 |  |  |  |  |  |
| A475:S31 | -0.15 | -0.44 | -0.22 |  |  |  |  |  |
| A475:S105 | 0.59 | 0.23 | 0.23 |  |  |  |  |  |
| A475:C106 | 0.36 | 0.37 | 0.31 |  |  |  |  |  |
| A475:D108 | 0.27 | 0.25 | 0.21 |  |  |  |  |  |
| G476:S105 | -0.13 | -0.36 | -0.33 |  |  |  |  |  |
| G476:C106 | -0.14 | -0.28 | -0.23 |  |  |  |  |  |
| G476:D108 | 2.48 | 2.38 | 1.83 |  |  |  |  |  |
| S477N:S105 | -0.95 | -1.43 | -1.15 |  |  |  |  |  |
| S477N:C106 | -0.44 | -0.74 | -0.48 |  |  |  |  |  |
| S477N:N107 | -0.16 | -0.29 | -0.25 |  |  |  |  |  |
| S477N:D108 | 6.42 | 5.61 | 5.46 |  |  |  |  |  |
| T478K:D108 | 3.32 | 3.85 | 3.54 |  |  |  |  |  |
| T478K:S32* | -1.43 | -1.85 | -2.63 |  |  |  |  |  |
| T478K:Y33* | 0.83 | -0.41 | -0.24 |  |  |  |  |  |
| P479:S32* | 0.32 | 1.04 | 0.93 |  |  |  |  |  |
| P479:Y33* | 1.71 | 2.30 | 2.15 |  |  |  |  |  |
| N481:Y33* | 0.28 | 0.23 | 0.15 |  |  |  |  |  |
| V483:S94* | -0.38 | -0.18 | -0.65 |  |  |  |  |  |
| E484A:V52 | 0.17 | 0.17 | 0.17 |  |  |  |  |  |
| E484A:S55 | 0.43 | 0.43 | 0.43 |  |  |  |  |  |
| E484A:N57 | 2.22 | 2.23 | 2.23 |  |  |  |  |  |
| E484A:S94* | -0.53 | -0.20 | -1.06 |  |  |  |  |  |
| E484A:S95* | 0.02 | 0.33 | -0.15 |  |  |  |  |  |
| G485:S94* | -0.43 | -0.17 | -0.57 |  |  |  |  |  |
| F486V:D99 |  |  | 0.32 |  |  |  |  |  |
| F486V:C106 |  |  | 0.18 |  |  |  |  |  |
| F486V:N107 |  |  | 0.26 |  |  |  |  |  |
| F486V:D108 |  |  | 0.57 |  |  |  |  |  |
| F486V:G109 |  |  | 0.58 |  |  |  |  |  |
| F486V:F110 |  |  | 0.21 |  |  |  |  |  |
| F486V:Y33* |  |  | 0.26 |  |  |  |  |  |
| F486V:Y92* |  |  | 0.84 |  |  |  |  |  |
| N487:P99 | -0.13 | -0.21 | -0.38 |  |  |  |  |  |
| N487:C106 | 0.02 | 0.31 | 0.19 |  |  |  |  |  |
| N487:D108 | 4.75 | 4.27 | 3.47 |  |  |  |  |  |
| Y489:S31 | 0.65 | 0.26 | 0.19 |  |  |  |  |  |
| Y489:V52 | -0.19 | -0.21 | -0.20 |  |  |  |  |  |
| Y489:C106 | -0.21 | -0.31 | -0.30 |  |  |  |  |  |

| **Table S4: COV2-2130** | | | |
| --- | --- | --- | --- |
| RBD:mAb pair | ΔBA.1 | ΔBA.2 | ΔBA.4 |
| T345:D31 | -0.25 | -0.30 | -0.27 |
| T345:D56 | -0.21 | -0.25 | -0.18 |
| T345:T108 | -1.29 | -1.49 | -1.22 |
| R346:R30 | -0.72 | -0.73 | -0.73 |
| R346:D31 | 7.47 | 7.51 | 7.49 |
| R346:W33 | -0.79 | -0.77 | -0.70 |
| R346:S53 | 0.52 | 0.48 | 0.46 |
| R346:I55 | 1.78 | 1.58 | 1.32 |
| R346:Y106 | -5.01 | -4.92 | -4.72 |
| R346:D107 | -1.31 | -1.20 | -1.29 |
| R346:T108 | -0.56 | -0.25 | -0.63 |
| K444:Y104 | 0.30 | 0.68 | 0.56 |
| K444:V109 | -0.18 | -0.22 | -0.16 |
| V445:Y55* | 0.73 | 0.48 | 0.34 |
| V445:Y104 | 0.45 | 0.57 | 0.20 |
| V445:P111 | -0.19 | -0.16 | -0.14 |
| V445:G116 | 0.33 | 0.20 | 0.18 |
| G446S:K36* | 0.17 |  |  |
| G446S:Y55* | -0.98 |  |  |
| G446S:W56* | -1.85 |  |  |
| G446S:T59* | 0.35 |  |  |
| G446S:Y104 | -0.91 |  |  |
| G446S:Y105 | -0.64 |  |  |
| G446S:F118 | -0.69 |  |  |
| Y449:N34* | -0.27 | -0.27 | -0.21 |
| Y449:K36* | 0.68 | 0.28 | 0.37 |
| Y449:W56* | 0.22 | 0.03 | 0.07 |
| N450:Y106 | 0.40 | 0.23 | 0.30 |
| N450:D107 | 0.35 | 0.39 | 0.30 |
| L452R:Y31* |  |  | -1.37 |
| L452R:S33* |  |  | -2.68 |
| E484A:S32* | 2.35 | 2.19 | 2.38 |
| E484A:N35* | 0.34 | 0.31 | 0.34 |
| Q493R:S32* | -0.30 | -0.28 |  |
| Q493R:N34* | 0.31 | 0.31 |  |
| Q493R:N35* | -0.44 | -0.57 |  |
| Q498R:T59* | -0.10 | -0.28 | -0.25 |

| **Table S4: REGN10933** | | | | | | | | |
| --- | --- | --- | --- | --- | --- | --- | --- | --- |
| RBD:mAb pair | ΔBA.1 | ΔBA.2 | ΔBA.4 |  | RBD:mAb pair | ΔBA.1 | ΔBA.2 | ΔBA.4 |
| R403:T28 | 0.43 | 1.36 | 1.76 |  | Y489:Y33 | -0.18 | 1.49 | -0.08 |
| R403:S30 | -0.03 | -0.08 | -1.49 |  | Y489:Y50 | 0.98 | 1.01 | 0.97 |
| R403:D31 | -2.59 | -7.78 | -7.48 |  | Y489:Y53 | 1.12 | 1.15 | 0.25 |
| D405N:T28 |  | -1.03 | -0.77 |  | Y489:D99 | -5.14 | 0.16 | 0.13 |
| D405N:D31 |  | -0.17 | -0.20 |  | Y489:R100 | -0.77 | -1.20 | -0.26 |
| E406:T28 | 0.11 | -0.38 | -0.17 |  | Y489:T103 | -0.01 | -0.04 | -3.16 |
| E406:D31 | -0.66 | -0.20 | -0.19 |  | Y489:M104 | 0.58 | 0.36 | -0.13 |
| K417N:T28 | 0.32 | 0.32 | 0.30 |  | F490:Y33 | -0.25 | -0.02 | -0.07 |
| K417N:D31 | 12.34 | 12.36 | 11.86 |  | F490:Y53 | -0.15 | 0.19 | -2.56 |
| K417N:Y32 | 0.18 | 0.18 | 0.17 |  | P491:Y53 | 0.06 | 0.06 | -0.29 |
| K417N:T102 | 0.38 | 0.18 | 0.47 |  | L492:Y53 | -0.04 | -0.09 | -0.64 |
| Y421:T102 | 1.02 | 0.67 | 1.04 |  | Q493R:S30 | 3.20 | 3.14 |  |
| Y453:S30 | 1.43 | 1.41 | 1.00 |  | Q493R:D31 | -0.40 | -0.72 |  |
| Y453:D31 | 2.18 | 2.13 | 2.56 |  | Q493R:Y32 | -0.11 | 0.07 |  |
| L455:D31 | 0.43 | 0.42 | -0.42 |  | Q493R:Y33 | -1.65 | -1.08 |  |
| L455:Y53 | 2.04 | 2.04 | 0.34 |  | Q493R:Y53 | -1.60 | -1.91 |  |
| L455:G101 | -0.89 | -0.38 | 0.31 |  | Q493R:S54 | 1.38 | 1.38 |  |
| L455:T102 | 2.32 | 1.05 | 3.29 |  | Q493R:R72 | -0.29 | -0.32 |  |
| F456:Y53 | 0.74 | 0.74 | -0.21 |  | Q493R:R100 | -0.10 | -0.56 |  |
| F456:R100 | -0.86 | -0.52 | 0.08 |  | S494:Y53H | -0.83 | -1.46 | 0.00 |
| F456:G101 | -0.38 | -0.13 | 0.48 |  | G496S:A75 | 0.21 | 0.21 | 0.15 |
| F456:T102 | 1.32 | 0.47 | 1.06 |  | Q498R:D73 | -0.29 | -1.10 | -1.86 |
| F456:M104 | 1.27 | 1.40 | 1.98 |  | Q498R:A75 | 0.42 | 0.38 | 0.25 |
| Y473:T103 | -0.13 | -0.03 | -2.07 |  | Q498R:K76 | 0.32 | 0.32 | 0.27 |
| Y473:M104 | 0.17 | 0.73 | 1.43 |  | T500:A75 | -0.31 | -0.81 | -0.24 |
| A475:R100 | 0.45 | 1.77 | 1.76 |  | N501Y:N74 | -0.37 | -1.51 | -0.49 |
| A475:M104 | -0.56 | -0.13 | -0.72 |  | N501Y:A75 | 0.09 | -0.54 | -1.09 |
| G476:Y32* | 0.42 | 0.38 | 0.44 |  | Y505H:G26 | 0.33 | 0.29 | 0.32 |
| G476:D92* | 0.72 | 0.71 | 0.63 |  | Y505H:F27 | 0.37 | 0.10 | 0.27 |
| G476:R100 | 0.15 | 0.26 | 0.24 |  | Y505H:N74 | 0.41 | 0.22 | 0.26 |
| S477N:T30* | -0.63 | -0.68 | -0.39 |  | Y505H:A75 | 0.99 | 0.82 | 0.39 |
| S477N:N31* | -1.38 | -1.86 | -1.52 |  | Y505H:S77 | 0.31 | 0.25 | 0.11 |
| S477N:Y32* | 0.58 | 0.47 | 0.32 |  |  |  |  |  |
| S477N:D92* | 5.12 | 5.09 | 4.65 |  |  |  |  |  |
| T478K:D28* | -0.20 | -0.56 | -0.59 |  |  |  |  |  |
| T478K:T30* | -1.00 | -1.73 | -0.21 |  |  |  |  |  |
| T478K:Y32* | -3.29 | -2.09 | -0.42 |  |  |  |  |  |
| T478K:D92* | -8.17 | -8.03 | -7.61 |  |  |  |  |  |
| E484A:Y33 | 0.41 | 0.47 | 0.44 |  |  |  |  |  |
| E484A:T52 | 2.44 | 2.44 | 2.42 |  |  |  |  |  |
| E484A:S56 | 0.86 | 0.86 | 0.85 |  |  |  |  |  |
| E484A:T57 | 2.99 | 3.01 | 2.93 |  |  |  |  |  |
| G485:Y33 | 0.35 | 0.41 | 0.32 |  |  |  |  |  |
| G485:T57 | 0.35 | 0.36 | 0.32 |  |  |  |  |  |
| G485:Y59 | 1.14 | 1.39 | 0.98 |  |  |  |  |  |
| F486V:Y91* |  |  | 0.31 |  |  |  |  |  |
| F486V:N93* |  |  | 0.73 |  |  |  |  |  |
| F486V:L94* |  |  | 0.61 |  |  |  |  |  |
| F486V:L96* |  |  | 0.93 |  |  |  |  |  |
| F486V:Y50 |  |  | 0.47 |  |  |  |  |  |
| F486V:Y59 | 1.65 | 1.48 | 1.62 |  |  |  |  |  |
| F486V:R100 |  |  | 0.32 |  |  |  |  |  |
| N487:Y32* | -1.45 | -2.60 | -0.41 |  |  |  |  |  |
| N487:Y91* | -1.32 | -0.80 | -0.87 |  |  |  |  |  |
| N487:N92* | 0.22 | -2.33 | -0.11 |  |  |  |  |  |
| N487:R100 | -4.07 | 1.32 | -3.51 |  |  |  |  |  |

| **Table S4: REGN10987** | | | | | | | | |
| --- | --- | --- | --- | --- | --- | --- | --- | --- |
| RBD:mAb pair | ΔBA.1 | ΔBA.2 | ΔBA.4 |  | RBD:mAb pair | ΔBA.1 | ΔBA.2 | ΔBA.4 |
| N343:Y102 | 0.02 | -1.42 | 0.02 |  | G446S:A33 | -0.45 |  |  |
| T345:T28 | 0.66 | 0.65 | 0.64 |  | G446S:Y35 | -0.25 |  |  |
| T345:N31 | 1.96 | 0.95 | 1.09 |  | G446S:W47 | 1.29 |  |  |
| T345:Y32 | 1.23 | 1.16 | 0.75 |  | G446S:I51 | -0.38 |  |  |
| R346:S30 | 0.92 | 0.72 | 0.82 |  | G446S:S52 | -1.11 |  |  |
| R346:N31 | 2.93 | 2.89 | 1.52 |  | G446S:Y53 | -0.25 |  |  |
| R346:Y53 | 3.75 | 1.48 | 1.85 |  | G446S:D54 | -0.15 |  |  |
| R346:D54 | 2.41 | -2.33 | 2.25 |  | G446S:S56 | -0.17 |  |  |
| R346:D101 | -2.90 | -0.02 | -0.03 |  | G446S:N57 | -1.83 |  |  |
| R346:Y102 | -1.02 | -0.01 | 0.00 |  | G446S:W99* | 2.00 |  |  |
| W436:Y102 | 0.01 | -1.57 | 0.01 |  | G447:V50 | 0.88 | 0.91 | 0.91 |
| N437:Y102 | 0.03 | -0.73 | 0.02 |  | G447:N57 | -0.92 | -0.04 | -1.29 |
| N437:Y32* | -1.36 | -0.01 | 0.00 |  | G447:Y59 | -0.23 | -0.80 | 0.19 |
| S438:Y102 | 0.03 | -0.73 | 0.04 |  | G447:W99* | 0.24 | 0.23 | 0.24 |
| N439:D104 | 2.06 | 4.31 | 0.15 |  | N448:A33 | 0.56 | 0.57 | 0.57 |
| N439:Y34* | -1.15 | -1.30 | -0.03 |  | N448:S52 | 0.23 | 0.22 | 0.21 |
| N440K:S100 | 0.47 | 0.43 | 0.35 |  | N448:Y53 | 1.48 | 1.62 | 1.37 |
| N440K:D101 | 0.08 | -0.13 | -1.03 |  | N448:N57 | -0.03 | -0.43 | -0.28 |
| N440K:Y102 | 1.84 | 0.65 | 0.19 |  | Y449:V50 | 0.33 | 0.33 | 0.33 |
| N440K:G103 | 0.80 | 0.52 | 0.34 |  | Y449:Y53 | -0.65 | 0.07 | 0.05 |
| N440K:D104 | -7.87 | -1.17 | -5.45 |  | Y449:S56 | -0.48 | 0.00 | -0.05 |
| N440K:N33* | -3.72 | -1.43 | 0.00 |  | Y449:N57 | 1.11 | 1.63 | 1.39 |
| N440K:Y34* | -2.46 | -2.44 | 0.07 |  | Y449:K58 | 0.20 | 0.20 | 0.21 |
| N440K:Y51* | 0.05 | -0.18 | -0.28 |  | Y449:Y59 | 1.84 | 1.70 | 1.85 |
| N440K:D52* | -7.89 | -7.15 | -7.14 |  | N450:S52 | 0.85 | 0.85 | 0.86 |
| L441:Y32 | 1.00 | 0.96 | 0.77 |  | N450:Y53 | 0.92 | 1.92 | 1.56 |
| L441:G99 | 0.17 | 0.16 | 0.16 |  | N450:D54 | 2.67 | 2.55 | 1.81 |
| L441:S100 | 0.45 | 0.35 | 0.33 |  | N450:N57 | 2.73 | 2.08 | 2.73 |
| L441:D101 | -0.04 | -0.30 | -0.48 |  | Y451:Y53 | 0.73 | 0.68 | 0.68 |
| L441:D104 | 0.07 | -1.07 | -0.10 |  | G496S:N57 | -0.64 |  |  |
| D442:Y53 | 0.37 | 0.32 | 0.26 |  | G496S:Y59 | -0.16 |  |  |
| S443:D104 | 5.44 | 6.26 | 6.07 |  | Q498R:Y59 | -3.48 | 0.00 | -0.22 |
| S443:Y105 | 0.10 | -1.36 | 0.09 |  | Q498R:D104 | 1.01 | 1.10 | 1.07 |
| K444:A33 | 2.44 | 2.63 | 2.64 |  | Q498R:S97* | 0.30 | -1.23 | 0.40 |
| K444:M34 | 0.58 | 0.59 | 0.59 |  | Q498R:W99* | 1.47 | 1.49 | 1.50 |
| K444:Y35 | 1.42 | 1.74 | 1.74 |  | P499:D104 | 0.61 | 1.75 | 0.97 |
| K444:V50 | 0.70 | 0.68 | 0.72 |  | P499:Y105 | 1.55 | 1.11 | 0.84 |
| K444:S52 | 0.02 | -0.50 | -0.79 |  | P499:Y34* | -0.51 | -1.33 | 0.14 |
| K444:Y53 | 0.00 | -0.08 | -2.36 |  | P499:L93* | 0.45 | -0.26 | 0.67 |
| K444:D54 | -0.05 | -0.66 | -7.80 |  | P499:S97* | 0.00 | -0.87 | -0.28 |
| K444:S56 | 0.00 | -0.52 | -0.87 |  | P499:W99* | 0.19 | 0.85 | 1.18 |
| K444:Y59 | -0.04 | -1.95 | -0.05 |  | T500:Y105 | -0.01 | 0.03 | -0.85 |
| K444:S98 | 3.61 | 3.60 | 3.59 |  | T500:Y32* | 1.78 | -0.77 | 1.14 |
| K444:G99 | 0.84 | 1.06 | 1.04 |  | T500:L93* | 1.07 | 0.47 | 0.25 |
| K444:S100 | -0.13 | 0.47 | 0.47 |  | T500:T94* | 0.24 | 0.00 | -1.55 |
| K444:D101 | -1.73 | 0.08 | 0.06 |  | T500:S95* | 0.03 | -0.78 | -0.34 |
| K444:D104 | 1.35 | 1.27 | 2.97 |  | T500:S97* | -0.19 | -0.67 | -0.10 |
| K444:Y105 | 0.23 | -0.48 | 0.45 |  | T500:W99* | 0.21 | 0.80 | 0.33 |
| V445:Y35 | 1.17 | 2.36 | 2.15 |  | N501Y:L93* | -0.15 | 0.01 | -0.38 |
| V445:W47 | 2.56 | 2.37 | 2.84 |  | N501Y:S95* | -0.01 | -0.02 | -1.02 |
| V445:S52 | -0.32 | -0.01 | -0.85 |  | G502:L93* | -0.17 | 0.00 | -0.32 |
| V445:N57 | -0.35 | -0.03 | -2.95 |  | G502:S97* | -0.49 | 0.00 | 0.00 |
| V445:Y59 | -1.19 | -2.48 | -1.48 |  | V503:Y32* | -0.20 | -0.14 | -0.67 |
| V445:W99* | 2.33 | 1.78 | 2.51 |  | V503:L93* | -0.32 | 0.00 | -0.29 |
| V445:D104 | 0.22 | 1.52 | 1.49 |  | Q506:Y34* | -1.37 | -0.15 | -0.06 |
| V445:Y105 | 1.08 | 1.63 | 2.10 |  | Q506:L93* | -0.31 | 0.00 | -0.31 |
| V445:S97* | -0.04 | -2.83 | -0.38 |  | R509:Y102 | -0.05 | -0.87 | 0.01 |

| **Table S4: S309** | | | |
| --- | --- | --- | --- |
| RBD:mAb pair | ΔBA.1 | ΔBA.2 | ΔBA.4 |
| N334:P28 | -0.33 | -0.38 | -0.34 |
| N334:T30 | 1.32 | 2.16 | 2.05 |
| N334:Y54 | 1.14 | 1.48 | 1.69 |
| N334:W105 | 0.61 | 1.06 | 1.04 |
| L335:P28 | 0.49 | 0.86 | 0.41 |
| L335:T30 | 0.19 | 0.20 | 0.17 |
| L335:S31 | 0.11 | 0.66 | 0.57 |
| L335:W105 | 0.68 | 0.71 | 0.75 |
| C336:W105 | 0.42 | 0.49 | 0.38 |
| P337:W105 | 0.31 | 0.28 | 0.25 |
| P337:F106 | 0.18 | 0.40 | 0.44 |
| G339D:Y32 | -4.22 | -0.83 | -2.62 |
| G339D:Y100 | -3.04 | -3.22 | -4.35 |
| G339D:L110 | -0.50 | -0.75 | -0.63 |
| E340:Y100 | 0.22 | 0.29 | 0.18 |
| E340:F106 | -0.90 | -0.96 | -0.72 |
| E340:G107 | -0.41 | -0.90 | -0.28 |
| E340:E108 | -1.18 | -1.63 | -0.54 |
| E340:L110 | 0.25 | 0.24 | 0.15 |
| N343:Y100 | 0.57 | 0.42 | 1.02 |
| N343:L110 | 0.15 | 0.12 | 0.16 |
| T345:T32* | -0.91 | 0.18 | 0.10 |
| T345:S33* | 0.66 | 0.53 | 0.38 |
| T345:H92* | 0.36 | 0.16 | 0.22 |
| T345:D93* | 0.22 | 0.18 | 0.12 |
| A344:E108 | -0.30 | -0.30 | -0.18 |
| T345:S109 | -0.78 | -0.73 | -0.76 |
| T345:I111 | 0.31 | 0.20 | 0.20 |
| R346:D93* | 4.92 | 4.89 | 4.71 |
| R346:E108 | -0.30 | -1.83 | -0.54 |
| K356:F106 | -0.42 | -0.07 | -0.34 |
| K356:E108 | -2.36 | -1.76 | -1.59 |
| I358:W105 | 0.14 | -0.45 | -0.31 |
| N360:W105 | 0.21 | 0.14 | 0.23 |
| C361:W105 | 0.47 | 0.51 | 0.45 |
| N440K:S31* | 0.36 | 0.25 | 0.50 |
| N440K:S53* | -0.66 | -0.72 | -0.34 |
| N440K:S54* | -0.20 | -0.27 | -0.20 |
| N440K:S66* | -0.19 | -0.17 | -0.27 |
| L441:S31* | 0.22 | 0.07 | 0.44 |
| L441:T32* | 0.20 | 0.15 | 0.14 |
| L441:I111 | 0.25 | 0.05 | 0.20 |
| K444:T28* | 0.34 | 0.10 | 0.37 |
| V445:S68* | 0.43 | 0.05 | 0.44 |

| **Table S4: S2X259** | | | | | | | | |
| --- | --- | --- | --- | --- | --- | --- | --- | --- |
| RBD:mAb pair | ΔBA.1 | ΔBA.2 | ΔBA.4 |  | RBD:mAb pair | ΔBA.1 | ΔBA.2 | ΔBA.4 |
| Y369:I28 | 0.27 | 0.24 | 0.26 |  | D405N:Y33* |  | 2.22 | 2.36 |
| Y369:S55 | -1.17 | -0.28 | -0.22 |  | D405N:S95* |  | 4.24 | 4.15 |
| Y369:G106 | 0.21 | 0.18 | 0.19 |  | R408S:A31* |  | 0.84 | 0.68 |
| N370:M54 | 0.45 | 0.26 | 0.49 |  | R408S:G32* |  | 0.89 | 0.96 |
| N370:S55 | 0.33 | 0.03 | 0.64 |  | R408S:Y33* |  | 1.86 | 1.60 |
| N370:K74 | 1.27 | 1.17 | 1.25 |  | R408S:D34* |  | 6.09 | 6.03 |
| N370:W107 | 0.20 | 0.20 | 0.20 |  | R408S:D109 |  | -1.86 | -0.77 |
| S371L/F:S55 | 1.08 | 1.10 | 1.11 |  | R408S:D110 |  | 7.99 | 7.47 |
| S371L/F:W107 | 0.39 | 0.39 | 0.39 |  | G502:S95* | -0.03 | 0.35 | 0.28 |
| A372:S55 | 1.29 | 1.27 | 1.25 |  | G502:S96* | 0.19 | 0.23 | 0.20 |
| A372:G56 | 0.32 | 0.32 | 0.31 |  | G502:L97* | 0.15 | 0.24 | 0.11 |
| A372:M57 | 1.20 | 1.02 | 1.57 |  | V503:Y93* | -0.30 | -0.15 | -0.20 |
| S373P:I52 | -0.62 | -0.36 | -0.07 |  | G504:S95* | 0.45 | 0.32 | 0.20 |
| S373P:S55 | -0.33 | -0.48 | -0.72 |  | Q506:S98* | -0.23 | -0.28 | -0.15 |
| S373P:M57 | -1.06 | -1.04 | -1.01 |  |  |  |  |  |
| S373P:N59 | -0.90 | -0.22 | -0.02 |  |  |  |  |  |
| S373P:A58 | -0.49 | -0.15 | -0.02 |  |  |  |  |  |
| F374:I52 | -0.80 | -0.46 | -0.22 |  |  |  |  |  |
| F374:M57 | 0.63 | 0.46 | 0.21 |  |  |  |  |  |
| S375F:R50 | -0.05 | -0.35 | -0.26 |  |  |  |  |  |
| S375F:I52 | -0.33 | 0.05 | -0.23 |  |  |  |  |  |
| S375F:M57 | 0.56 | 0.56 | 0.15 |  |  |  |  |  |
| S375F:N59 | 0.21 | 0.38 | 0.43 |  |  |  |  |  |
| S375F:D109 | -0.23 | -0.46 | -0.48 |  |  |  |  |  |
| S375F:G108 | -0.43 | -0.55 | -0.42 |  |  |  |  |  |
| S375F:Y93* | -0.52 | -1.24 | -1.01 |  |  |  |  |  |
| S375F:P100* | -1.38 | -0.97 | -1.03 |  |  |  |  |  |
| S375F:W102* | -0.27 | -0.59 | -0.49 |  |  |  |  |  |
| T376A:W107 |  | 1.98 | 1.59 |  |  |  |  |  |
| T376A:G108 |  | 0.60 | 0.56 |  |  |  |  |  |
| T376A:D109 |  | 0.21 | 0.18 |  |  |  |  |  |
| F377:G106 | 0.00 | 0.17 | 0.15 |  |  |  |  |  |
| F377:W107 | -0.09 | -0.44 | -0.23 |  |  |  |  |  |
| K378:Y104 | 1.51 | -0.52 | -0.73 |  |  |  |  |  |
| K378:Y105 | -0.05 | 0.27 | 0.23 |  |  |  |  |  |
| K378:G106 | -0.02 | 0.28 | 0.30 |  |  |  |  |  |
| K378:W107 | 0.00 | 0.33 | 0.32 |  |  |  |  |  |
| K378:D109 | -0.56 | -6.20 | -5.86 |  |  |  |  |  |
| K378:D110 | 0.97 | -0.25 | -0.48 |  |  |  |  |  |
| K378:D111 | -0.25 | -0.15 | -0.23 |  |  |  |  |  |
| C379:Y104 | 0.64 | -0.23 | -0.31 |  |  |  |  |  |
| C379:Y105 | 0.67 | -0.42 | -0.40 |  |  |  |  |  |
| Y380:N103 | 0.59 | -0.10 | -0.31 |  |  |  |  |  |
| Y380:Y104 | 0.10 | -0.20 | -0.24 |  |  |  |  |  |
| V382:Y105 | 0.26 | 0.34 | 0.15 |  |  |  |  |  |
| S383:F29 | 0.55 | 0.19 | 0.21 |  |  |  |  |  |
| S383:Y105 | -0.15 | -0.47 | -0.31 |  |  |  |  |  |
| P384:F29 | -0.25 | -0.67 | -0.29 |  |  |  |  |  |
| P384:Y32 | -1.10 | -0.89 | -0.59 |  |  |  |  |  |
| T385:F29 | -0.46 | -0.95 | -0.52 |  |  |  |  |  |
| T385:N30 | -1.25 | -0.04 | -0.07 |  |  |  |  |  |
| T385:Y32 | -1.11 | -0.24 | -0.23 |  |  |  |  |  |
| T385:M54 | 0.24 | 0.26 | 0.02 |  |  |  |  |  |
| K386:F29 | -0.65 | -0.53 | -1.58 |  |  |  |  |  |
| L387:F29 | -0.04 | -0.40 | -0.59 |  |  |  |  |  |
| L387:Y105 | 0.00 | -0.66 | -0.60 |  |  |  |  |  |

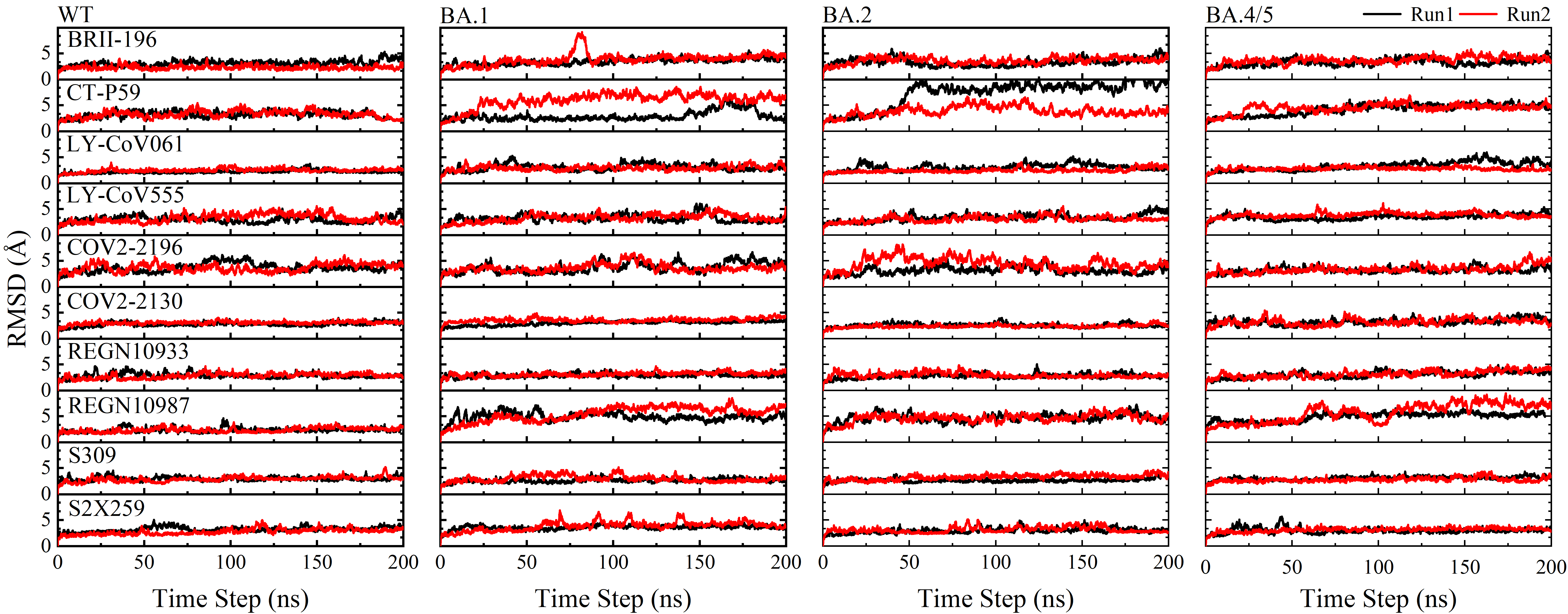

**Figure S1.** Time revolution of the root mean square deviation (RMSD) of the heavy atoms of the mAb-RBD complex in the WT and Omicron cases for both MD runs. All analyses are based on 5000 snapshots taken from each MD run over the course of 200 ns.

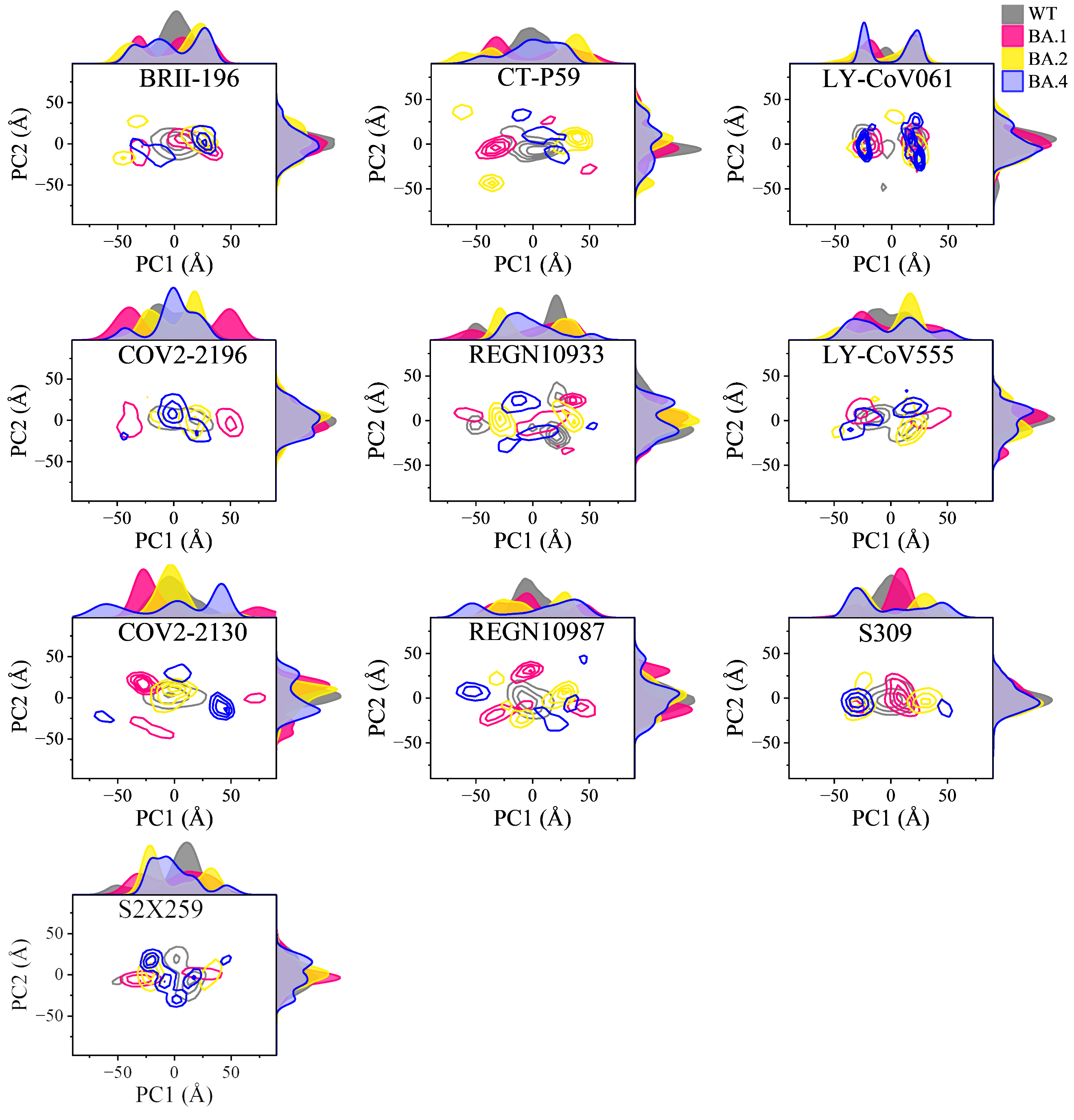

**Figure S2.** Dynamic conformations projected of two principal components (PC1 and PC2) of the RBD residues (T333-P527) for all mAb-RBD complexes in both WT and Omicron cases. Certain Omicron RBDs have more than one local minimums.

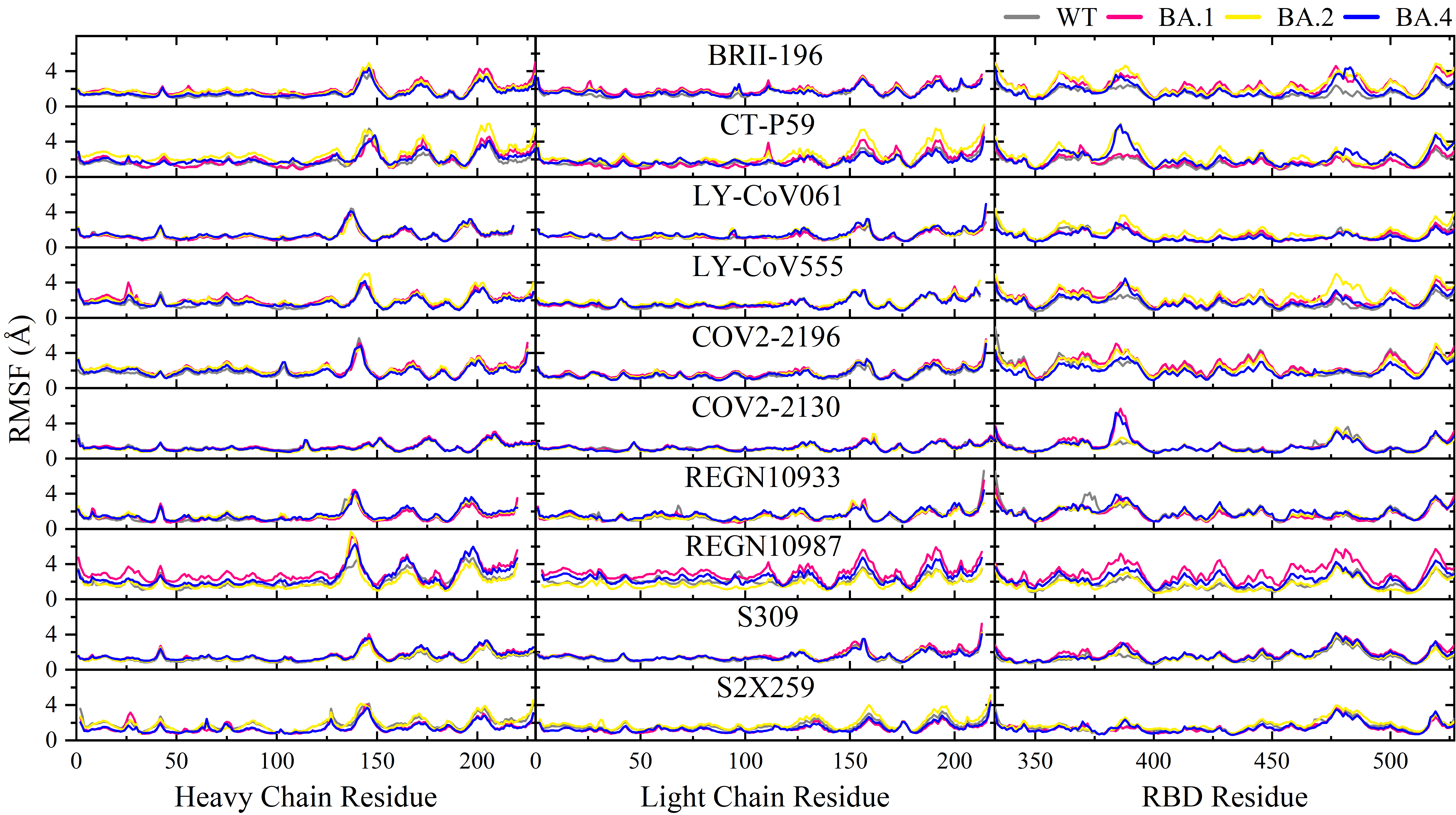

**Figure S3.** The root mean square fluctuation (RMSF) of the residues at mAb-RBD complex in all Omicron and WT cases. Heavy and light chain residues of mAb is the left and middle panels while RBD in right.

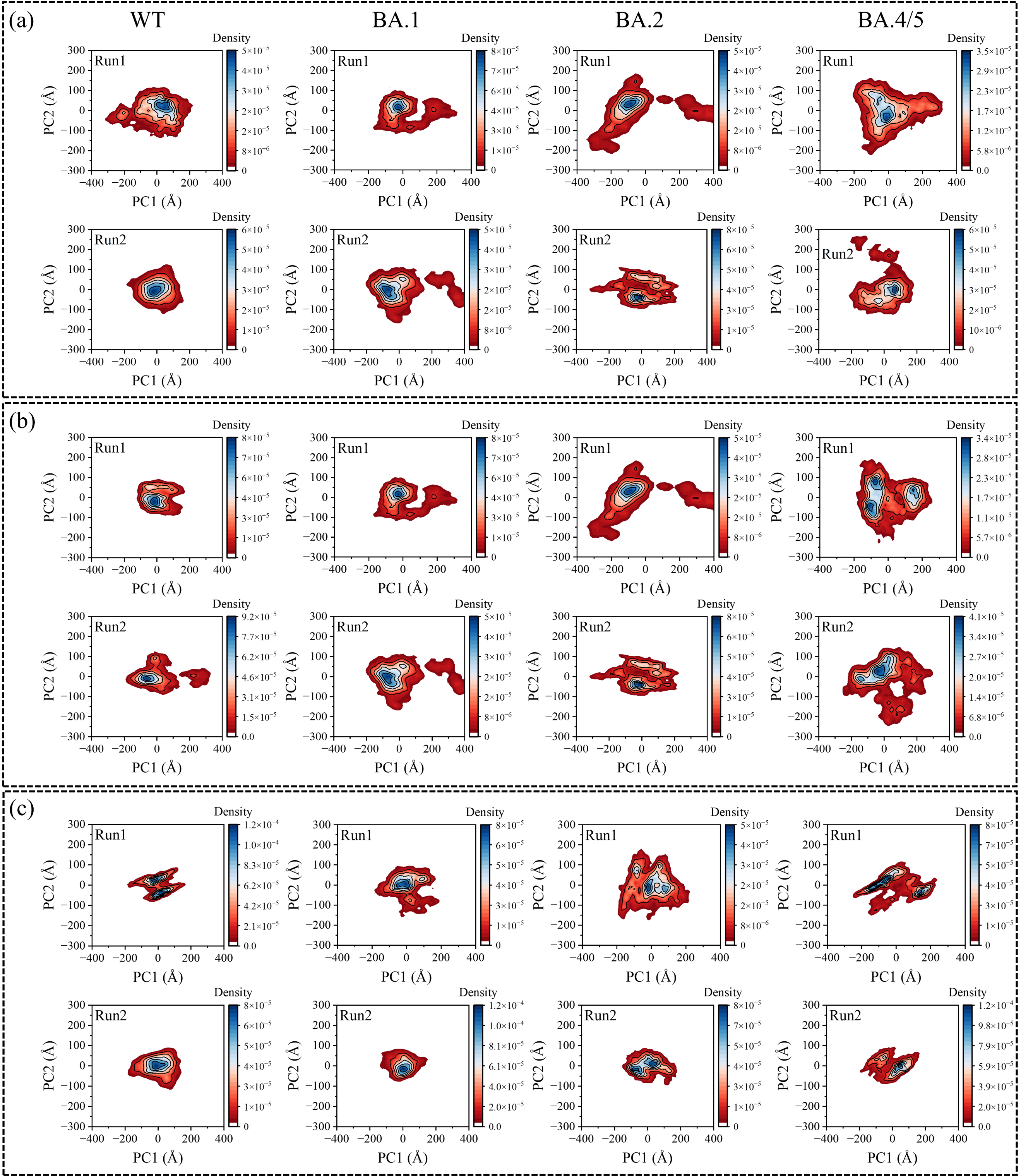

**Figure S4.** Dynamic conformations projected of two principal components (PC1 and PC2) of the class 1 mAb-RBD complexes in the WT and Omicron cases for both MD runs. The black lines are used to separate the main conformational dominants. (a) For BRII-196, (b) CT-P59 and (c) LY-CoV061.

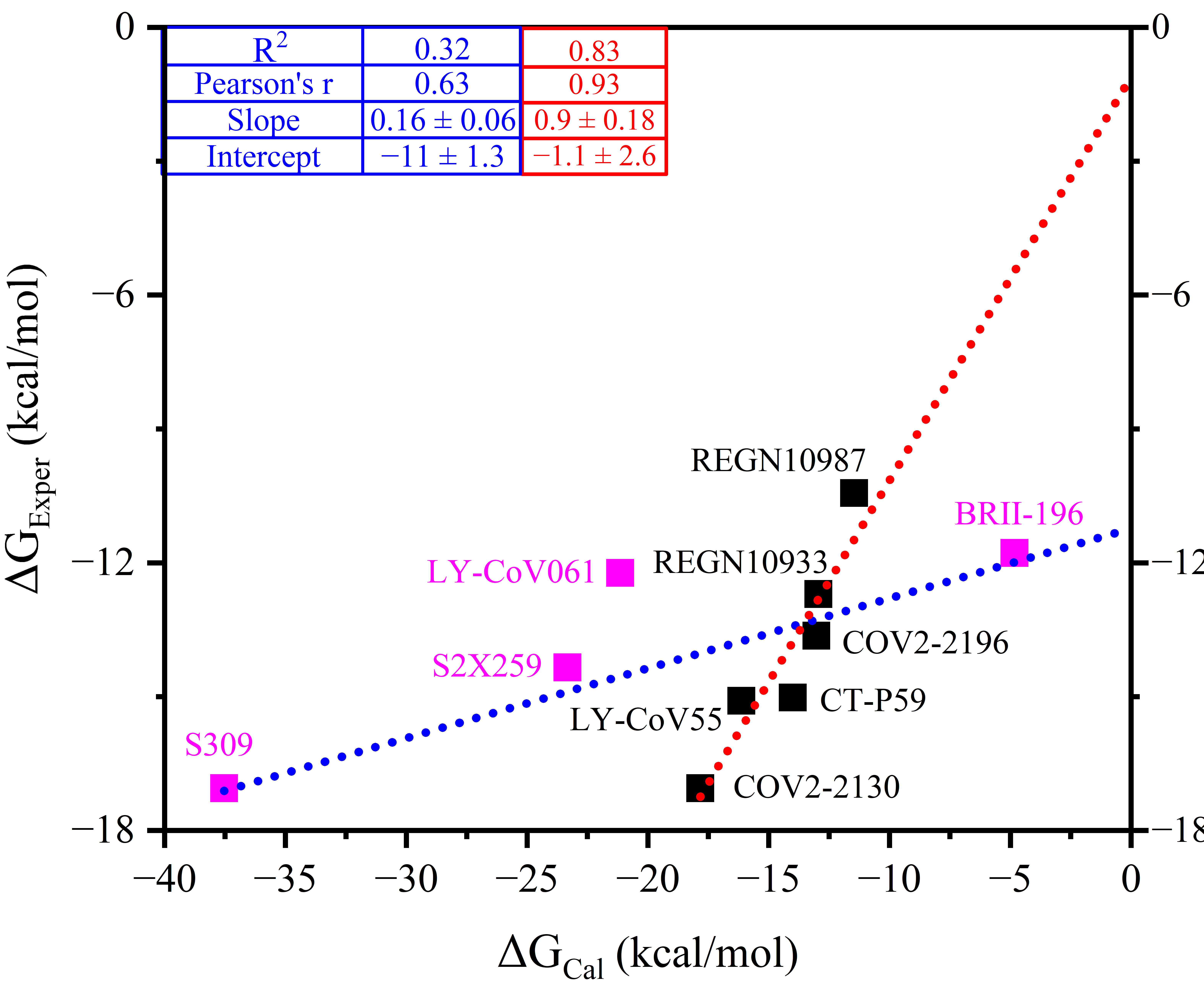

**Figure S5.** Scatter correlation plots of calculated versus experimental binding free energies for WT mAb-RBD complexes (see Table S2). The blue line is used to fit all 10 biosystems with their experimental values while the red line is for only six biosystems by excluding BRII-196, LY-CoV061, S309, and S2X259 (pink symbols).

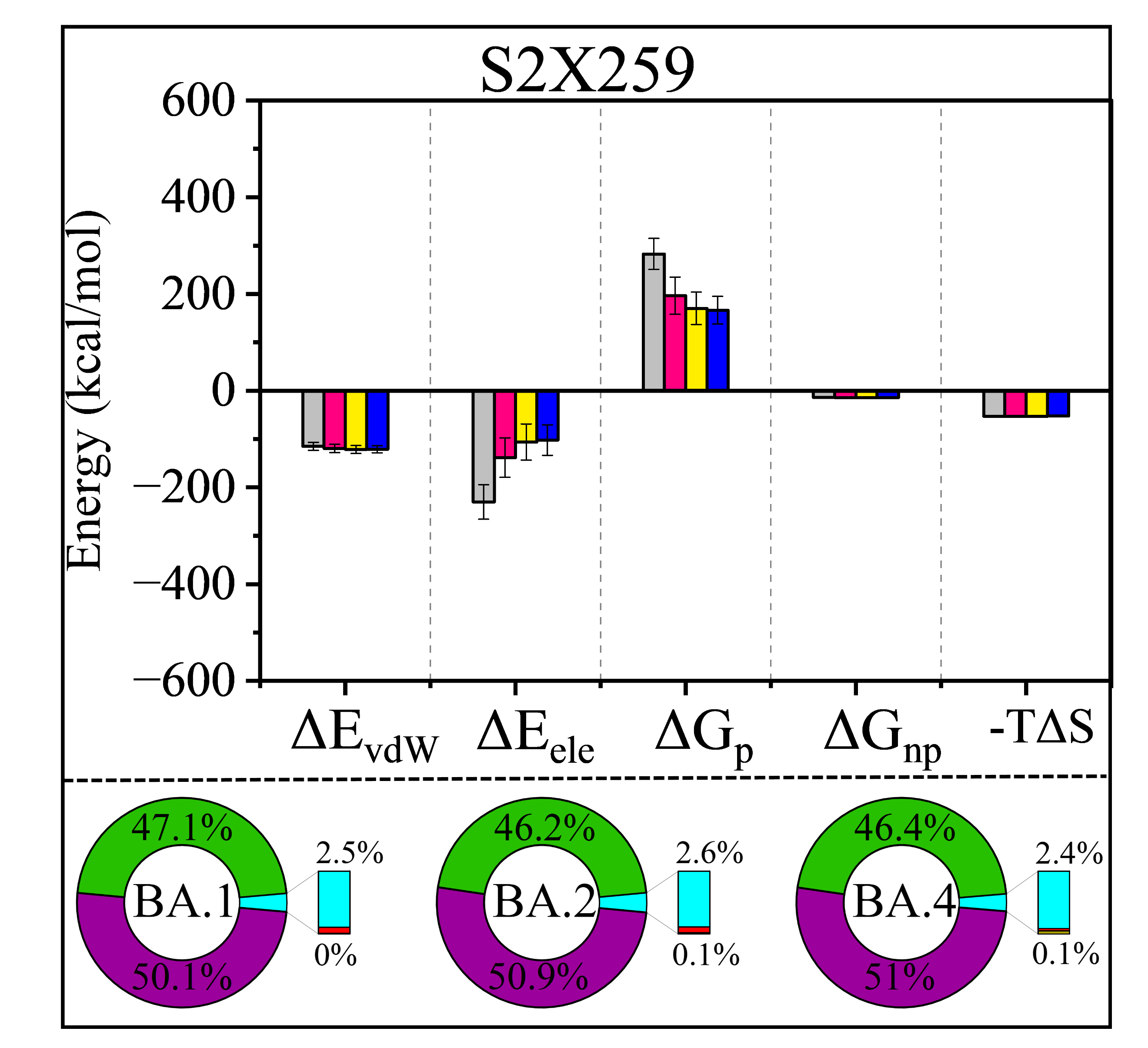

**Figure S6.** The electrostatic interaction role of Omicron subvariant mutations in conferring mAb resistant for S2X259. The histogram displays the BFE dissection in terms of energetic components including van der Waals (ΔE_vdW_), Coulombic electrostatic (ΔE_ele_), electrostatic polar (ΔG_p_), non-polar (ΔG_np_), and entropic (-TΔS) contributions. Because the standard error of mean (SEM) is too small to appear in this histogram, the standard deviation (SD) is reported instead. The doughnut plots in the bottom of each panel represent the relative energetic differences of Omicron subvariants with respect to its WT value.

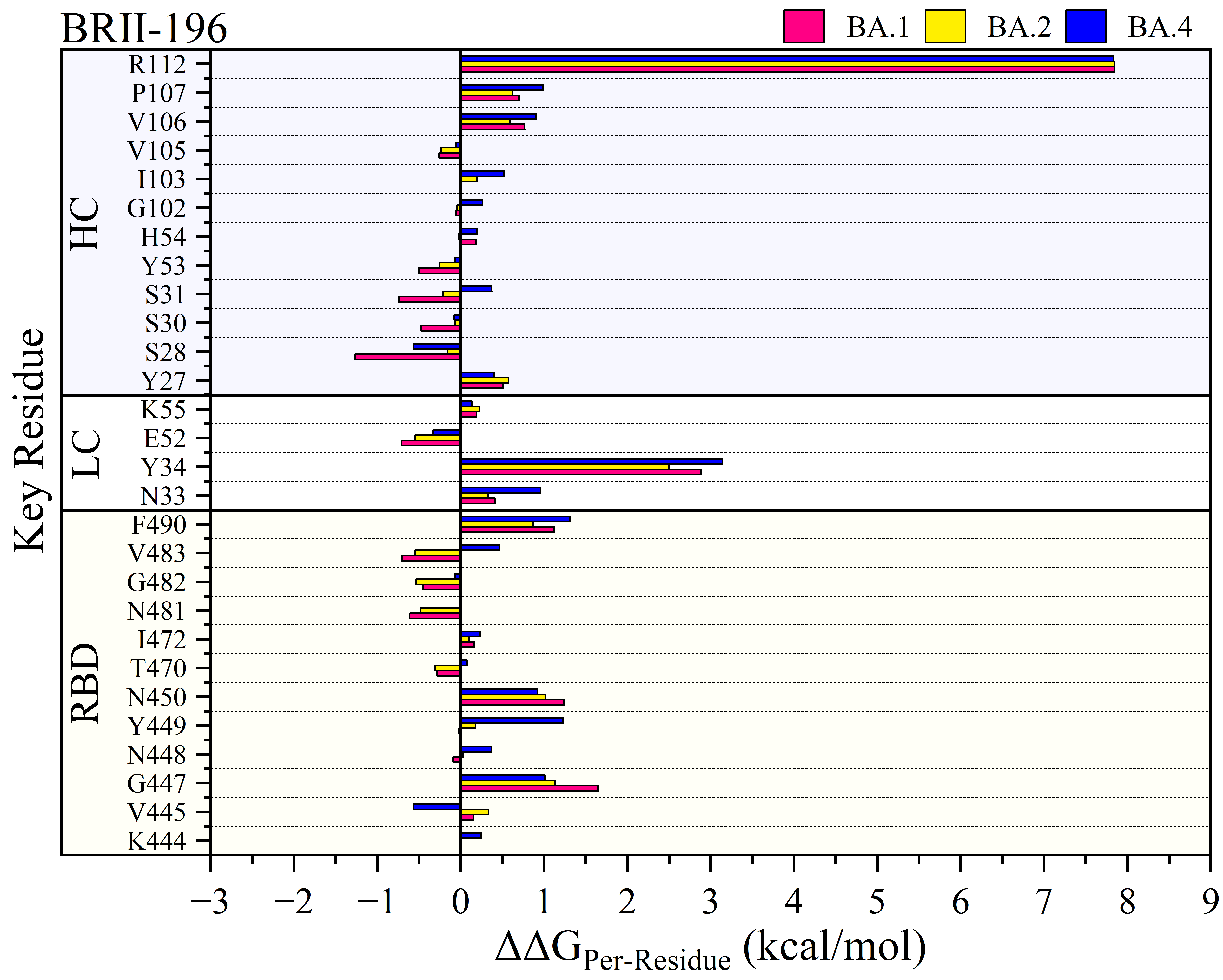

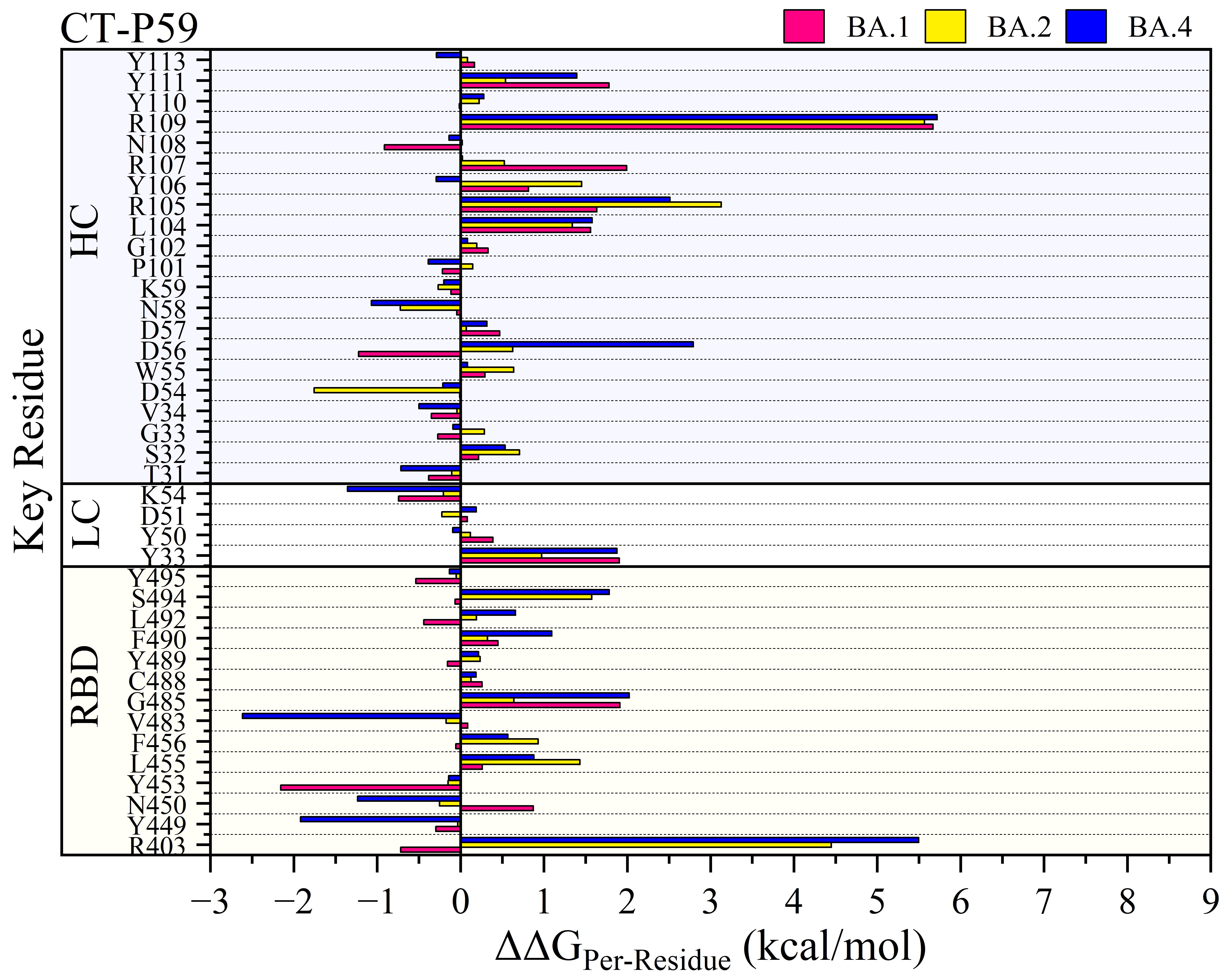

Continue
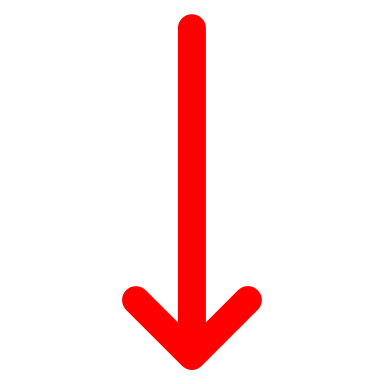

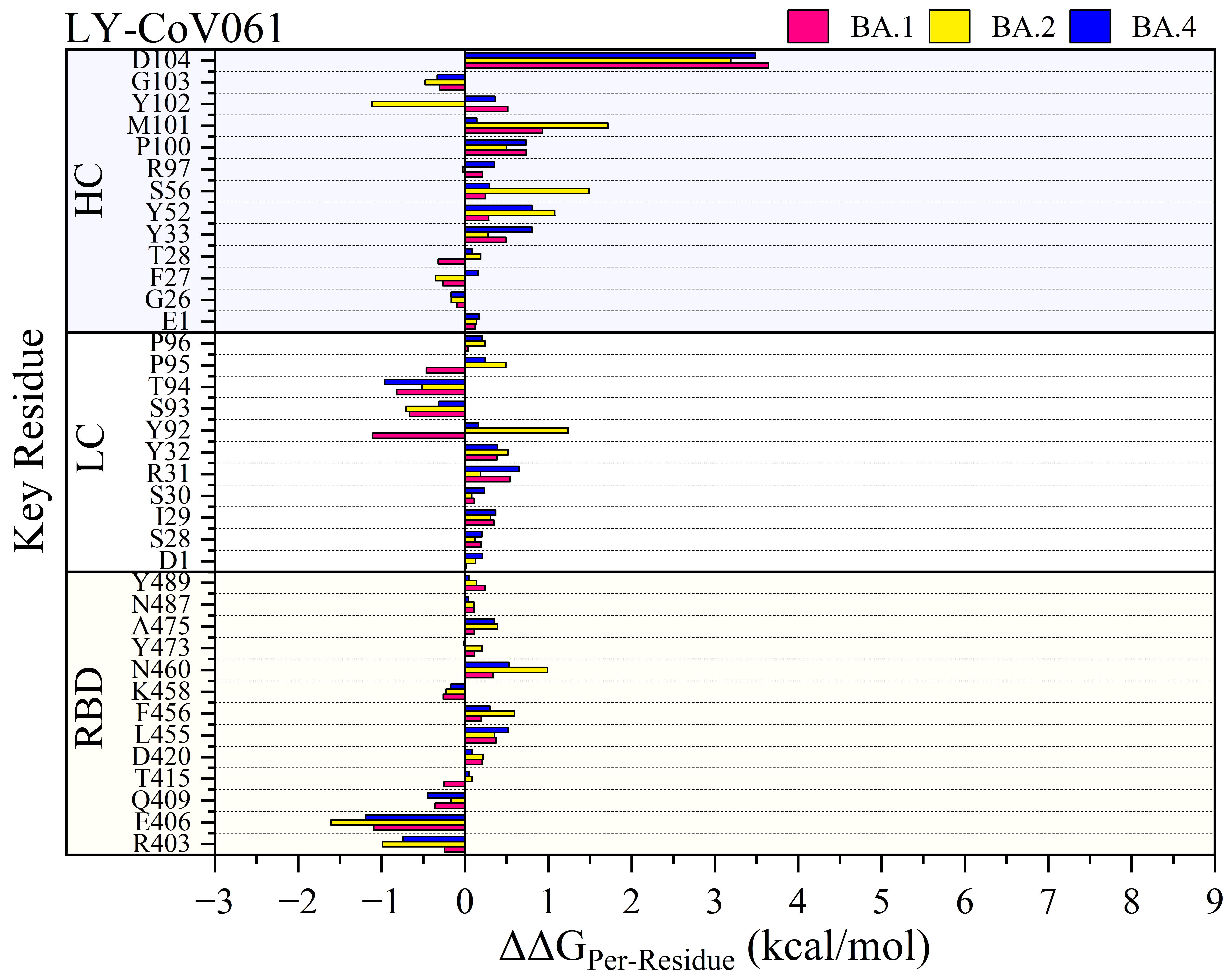

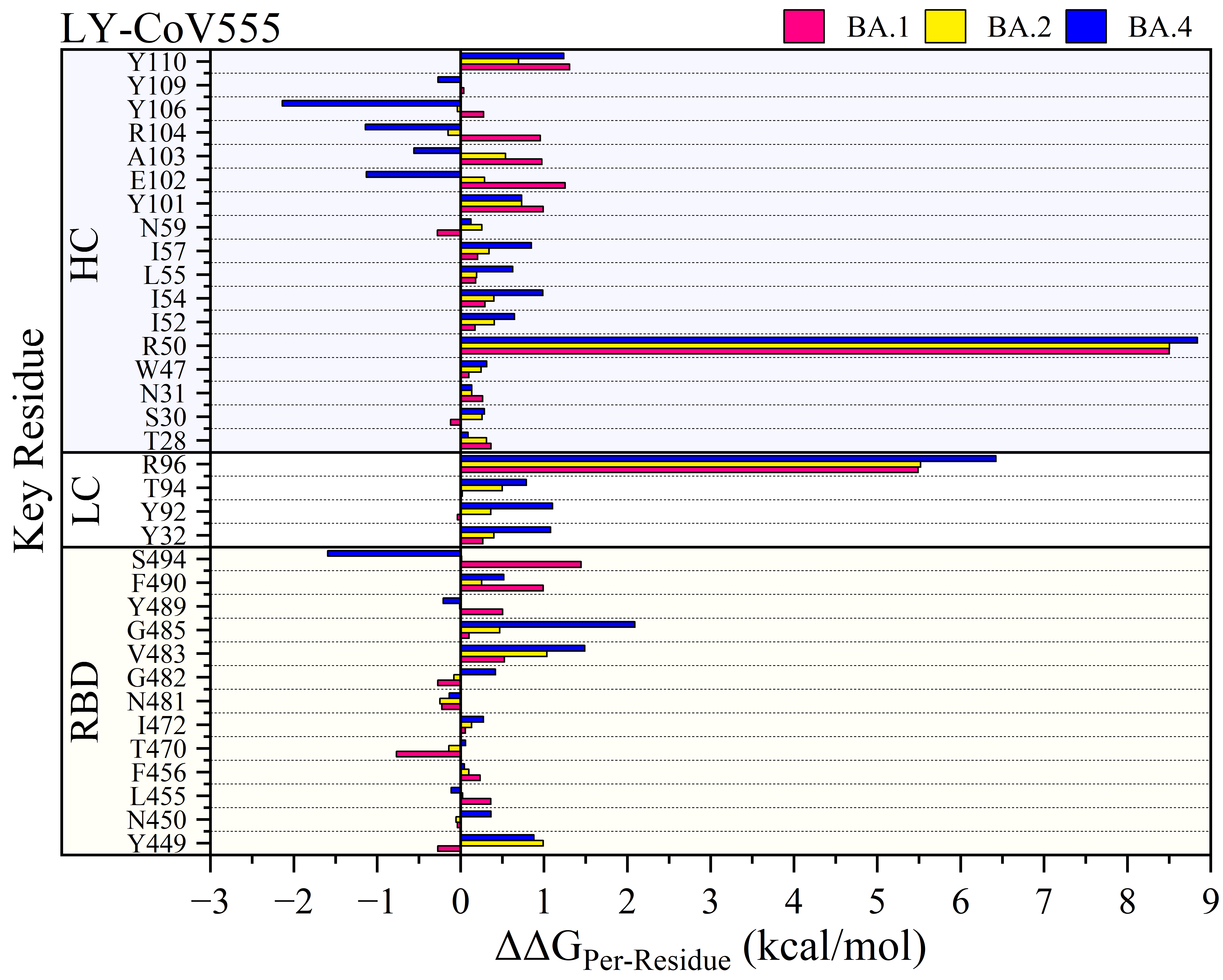

Continue
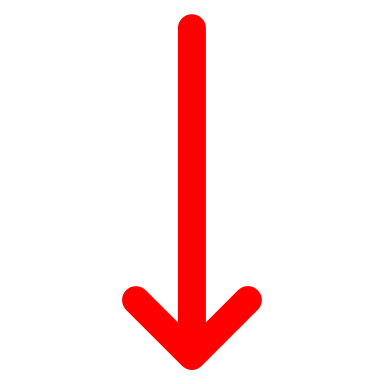

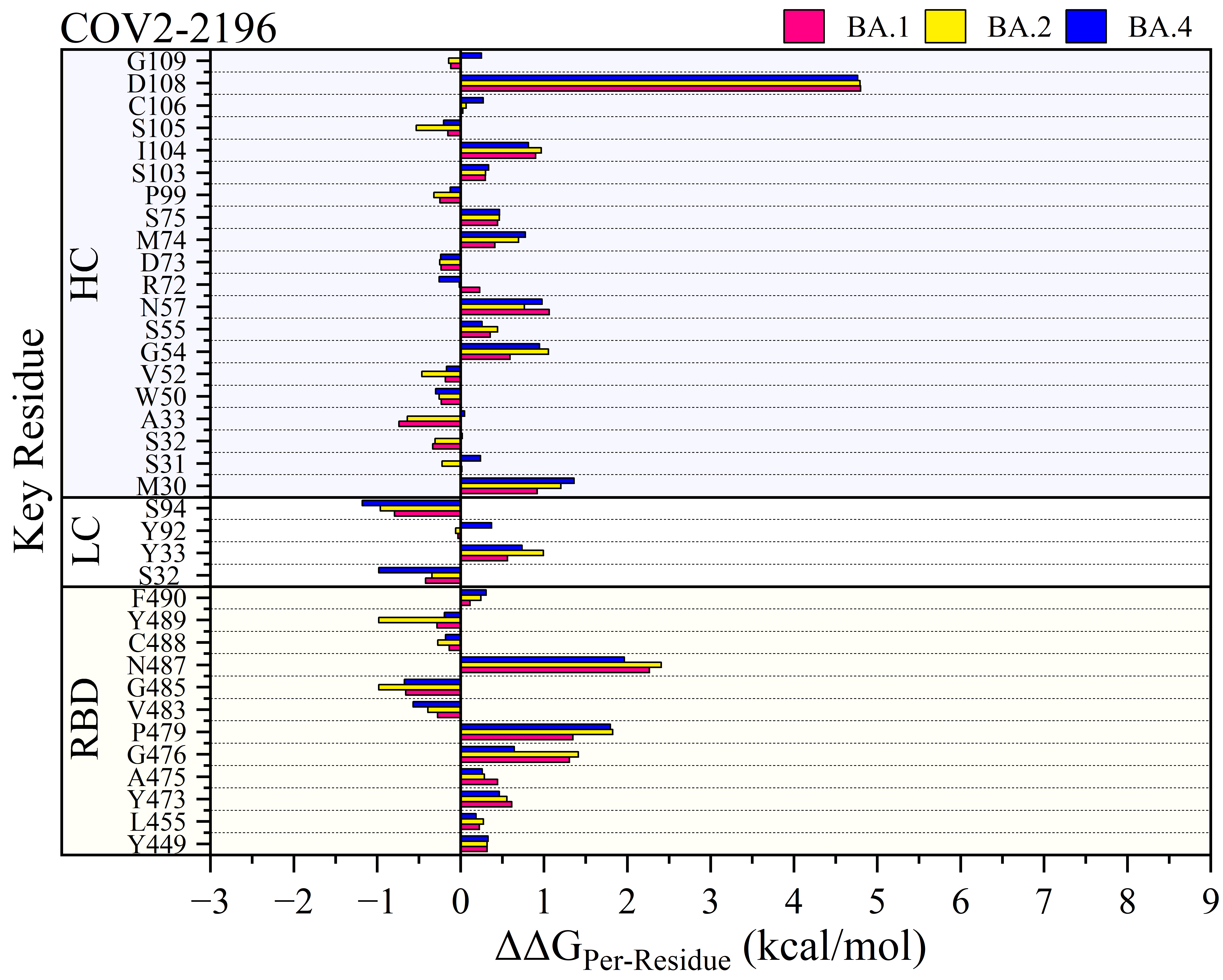

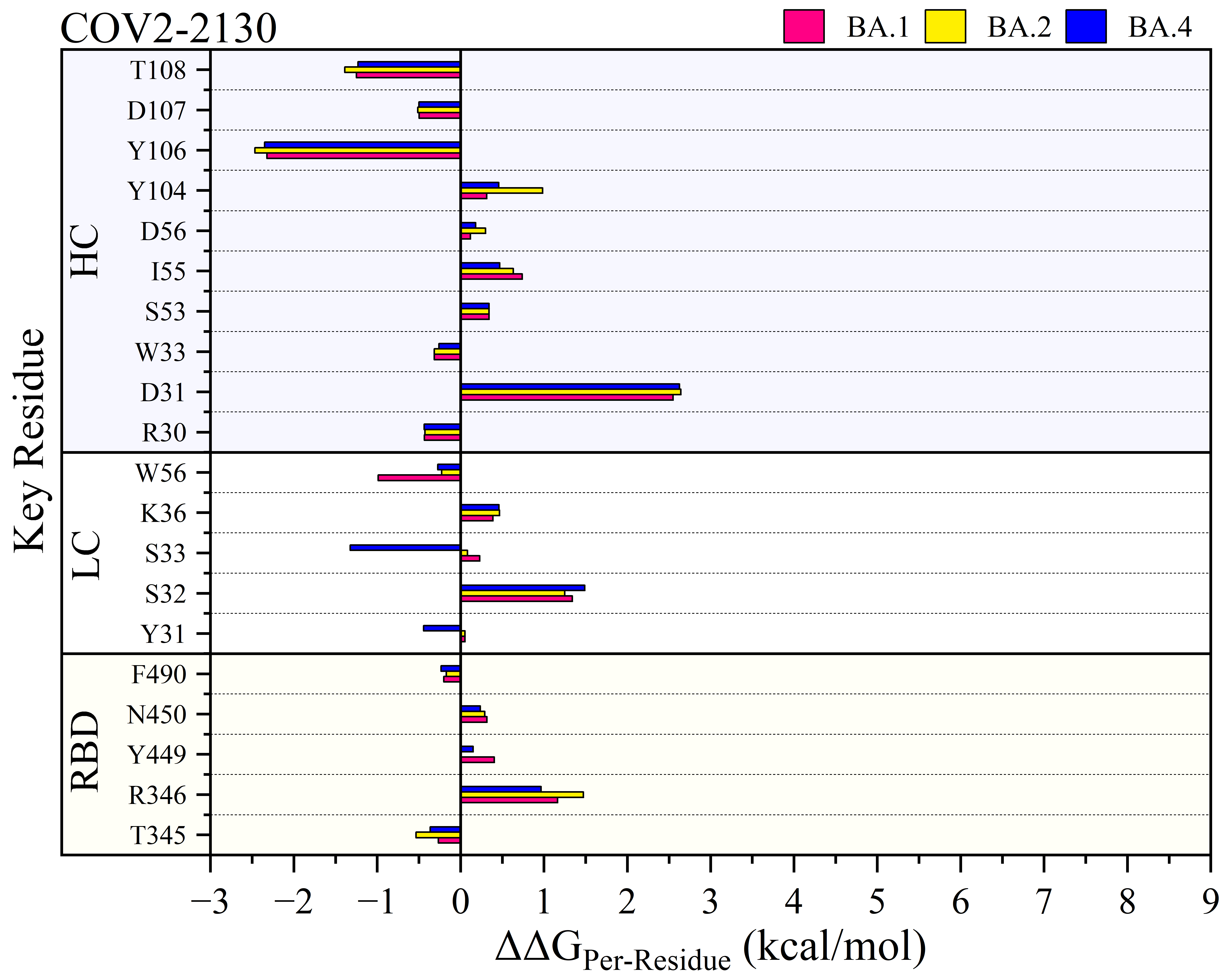

Continue
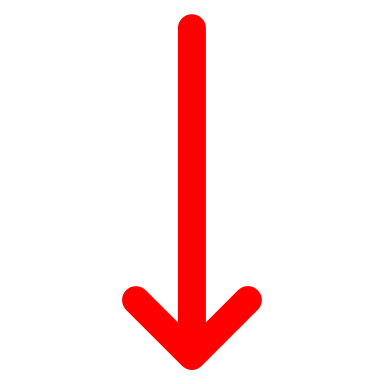

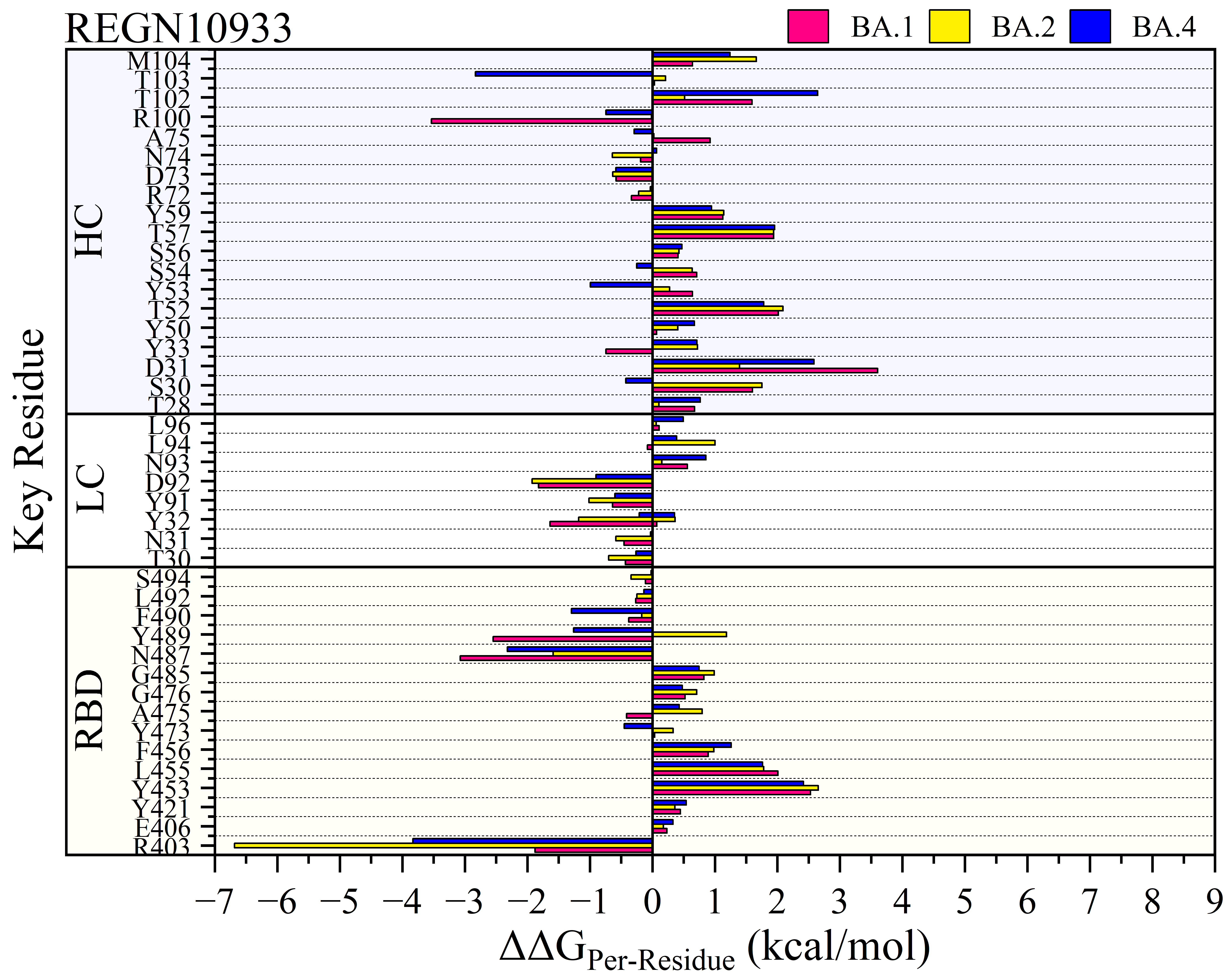

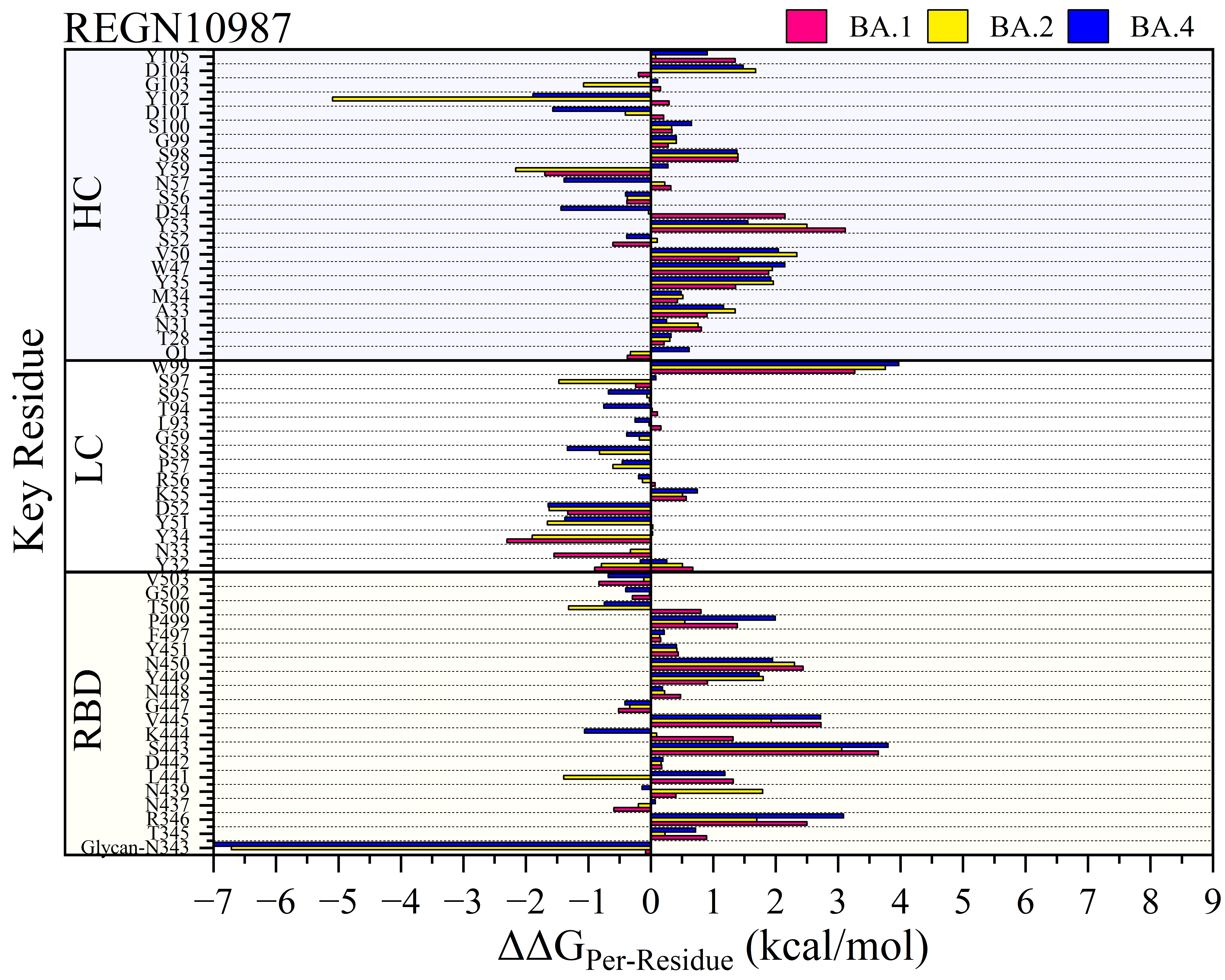

Continue
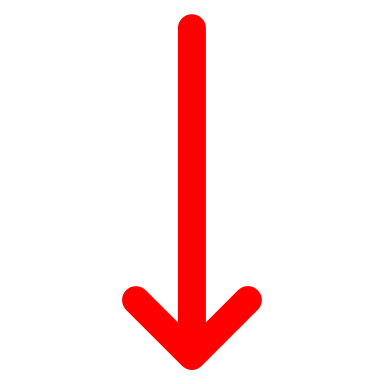

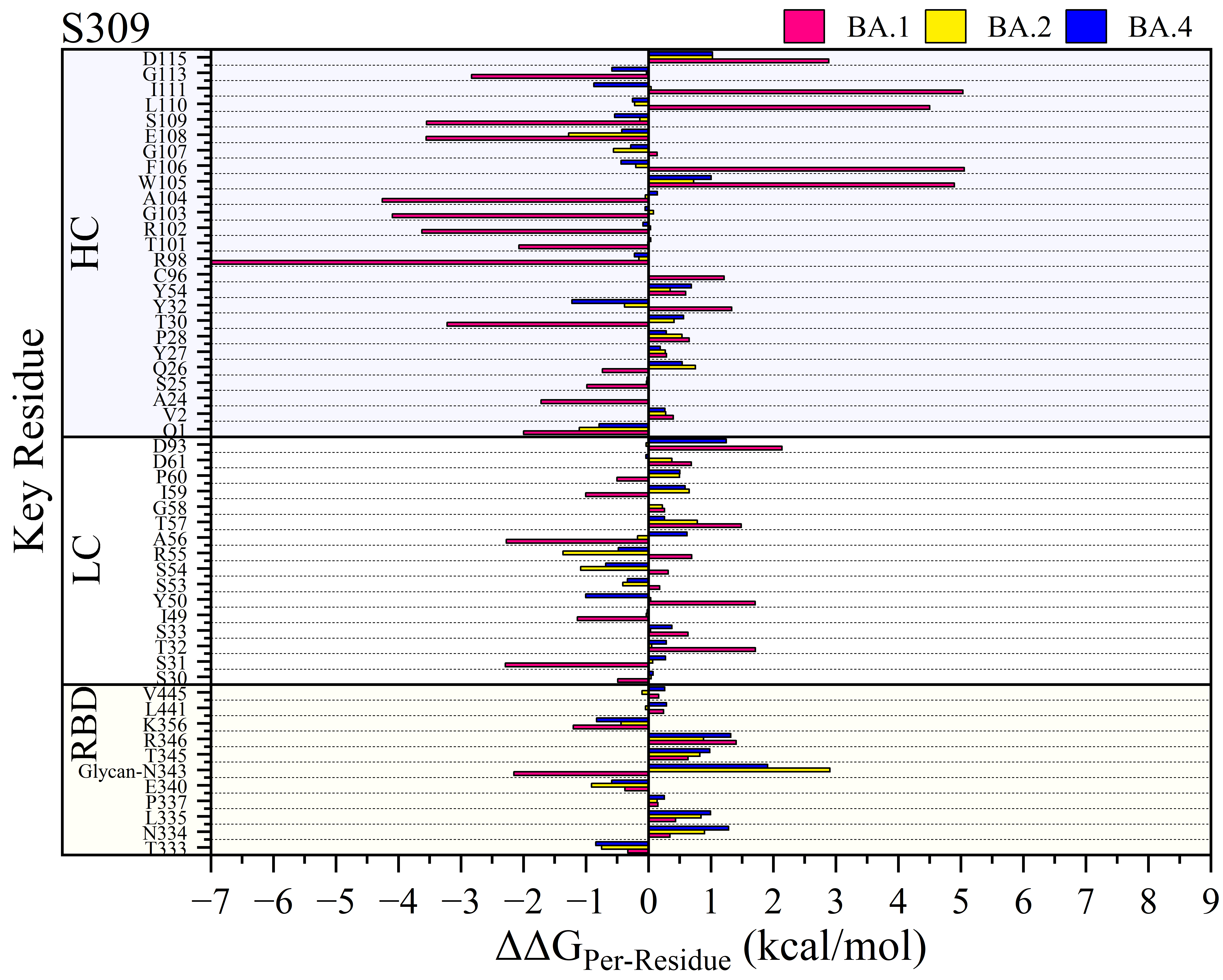

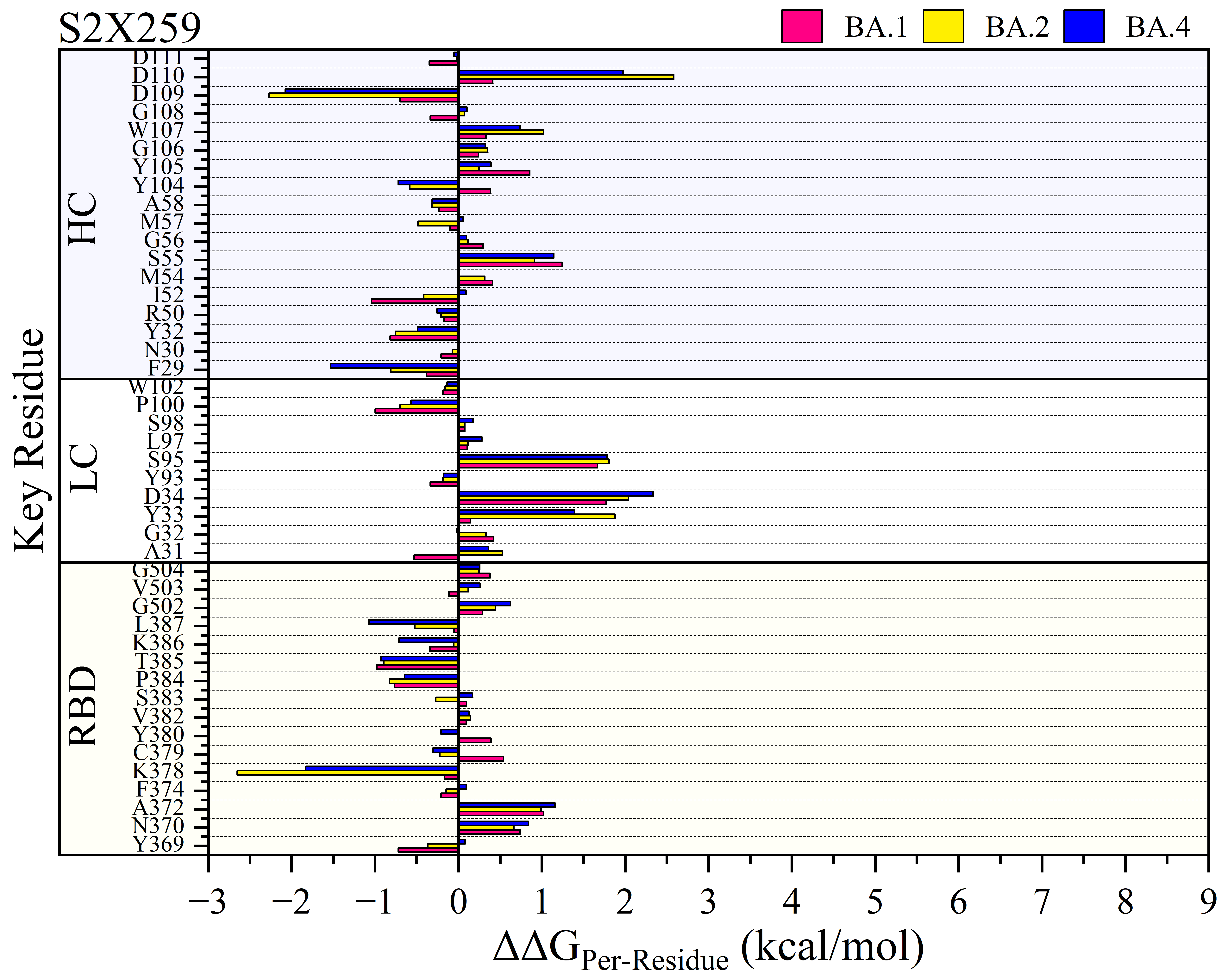

**Figure S7.** RBD Omicron mutations impact the interaction spectrum for all mAbs based on the per-residue energy decomposition of the key residues for both conserved RBD and heavy and light chains (HC or LC) of mAb. These analyses are based on the relative per-residue BFE (ΔΔG_per-residue_ in kcal/mol unit) of each RBD, HC or LC sites with respect to its WT. We ignore thermal fluctuation interactions by not including energy differences range between 0.15 to -0.15 kcal/mol.
